# Synthetic engineering demonstrates that synergy among enhancers involves an increase in transcriptionally productive enhancer-gene contacts

**DOI:** 10.1101/2025.08.08.669284

**Authors:** Endika Haro, Marianna Iliadou, Shyam Ramasamy, Thais Ealo, Victor M. Campa, Tomas Pachano, A. Marieke Oudelaar, Alvaro Rada-Iglesias

## Abstract

Enhancers are non-coding *cis*-regulatory elements that control the expression of distally located genes in a tissue- and time-specific manner. Recent studies indicate that enhancers can differ in their underlying genetic architecture and regulatory properties. However, these different types of enhancers were previously investigated under rather variable conditions (e.g. model organism, cell type, enhancer-promoter distance, type of target promoter, *etc*.), thus introducing confounding factors that make it difficult to discern the distinct regulatory properties of each enhancer type. To overcome these limitations, here we generated transgenic mouse embryonic stem cells (mESC) lines in which different types of synthetic enhancers (i.e. “typical” enhancer, CTCF-associated enhancer, enhancer cluster/super-enhancer) were built upon the same “core” neural enhancer and inserted at the same distance (i.e. 100 Kb) from a typical developmental gene (i.e. *Gata6*). Subsequently, the mESC lines were differentiated to systematically compare the regulatory properties of the different enhancer types under identical conditions.

Regarding the CTCF-associated enhancer, our data revealed that the addition of a CTCF site to the “core” enhancer increased insulation and led to the formation of a smaller contact domain, while having a rather mild effect on enhancer-promoter contact frequency and target gene expression. On the other hand, in comparison to the “core” enhancer alone, the enhancer cluster synergistically increased target gene expression and burst fraction. Importantly, we found that, in contrast to previous models, the strong regulatory activity of the enhancer cluster can not be explained by changes in enhancer-promoter contact frequency or the formation of transcriptional condensates. Instead, our data suggest that the emergent regulatory properties of enhancer clustering preferentially entail an increase in RNA Polymerase II pause release and, thus, in the fraction of enhancer-promoter contacts that are transcriptionally productive.

## Introduction

Enhancers are a diverse group of *cis*-regulatory elements that are able to regulate the expression of their target genes over long genomic distances (i.e. long-range gene regulation) in a time- and tissue-specific manner (Bower and Kvon, 2025; Noonan and McCallion, 2010). This long-range regulatory capacity is particularly prominent for enhancers controlling the expression of major developmental genes (e.g. *SHH, SOX9, MYC, PITX1*), which can be located hundreds of Kb from each other (Kragesteen et al., 2018; Lettice et al., 2002; Long et al., 2020; Yanchus et al., 2022) and relies, at least partly, on the three-dimensional (3D) organization of the genome. The genome is organized into self-interacting topological associating domains (TADs) that often correlate with gene regulatory domains where enhancers and their target gene promoters come into physical proximity (Dixon et al., 2012; Dowen et al., 2014; Lieberman-Aiden et al., 2009; Phillips-Cremins et al., 2013; Rao et al., 2014; Tang et al., 2015). These TADs might confer a permissive topological framework within which, according to the classical looping model, enhancers and gene promoters come into physical proximity (Long et al., 2016). In the case of long-range enhancers, the physical communication with the target promoters might require aid from tethering elements such as intra-TAD CTCF sites, CpG islands or LDB1-bound regions (Aboreden et al., 2025; Cruz-Molina et al., 2017; Kubo et al., 2021; Pachano et al., 2021). However, although enhancer-promoter contacts might be instructive for gene transcription in certain cellular or developmental contexts (Chen et al., 2024; Pollex et al., 2024), the overall picture is that proximity is neither sufficient for gene expression nor does it show a linear relationship with gene expression levels (Zuin et al., 2022). Therefore, alternative models of enhancer function that aim to reconcile the rather fuzzy correlation between enhancer-promoter contacts and gene transcription have been proposed (e.g. the transcriptional condensate/hub model or the TF activity gradient (TAG) model) (Boija et al., 2018; Karr et al., 2022; Sabari et al., 2018; Yang and Hansen, 2024). According to these alternative models, although certain level of physical proximity might be necessary for enhancer mediated transcriptional regulation, other factors and mechanisms (e.g. type and amount of co-activators) play a more instructive role in determining the levels of enhancer activity and target gene expression.

Transcription is a rather stochastic and discontinuous process that occurs in bursts separated by periods of inactivity (Bahar Halpern et al., 2015; Chubb et al., 2006). Therefore, the overall level of gene transcription can be modulated through two key parameters: burst frequency (i.e. the number of bursts in time units; inversely correlated with the OFF time) and burst size (i.e. amount of mRNA molecules produced in each burst; dependent on both burst amplitude and burst duration (ON time))(Tunnacliffe and Chubb, 2020). Previous studies indicate that burst size is primarily regulated at the promoter level, whereas burst frequency (or burst fraction, a related bursting parameter) is preferentially controlled by enhancers and the probability by which they contact with their target promoters (Bartman et al., 2016; Fukaya et al., 2016; Larson et al., 2013; Larsson et al., 2019; Walters et al., 1995). Recent live-cell imaging data suggest that enhancer-promoter contacts, which in general are rare, transient and highly dynamic (Chen et al., 2018; Mateo et al., 2019), can initiate bursts that are then maintained by promoters (hysteresis), thus potentially explaining the bursty nature of transcription and the relatively poor correlation between enhancer-promoter proximity and transcriptional output (Alexander et al., 2019; Xiao et al., 2021; Zuin et al., 2022). Furthermore, transitions in enhancer-promoter contacts seem to occur with faster dynamics than transcriptional bursts (Tünnermann et al., 2025), thus indicating that a significant fraction of such proximity events might not be transcriptionally productive. However, it is currently unknown which factors might determine whether an enhancer-promoter contact is productive or not.

In addition to the ongoing debate regarding how close enhancers and promoters need to be from each other to initiate a transcriptional burst, other fundamental aspects of enhancer function also remain unclear. Namely, which of the main stages of transcription (i.e. preinitiation complex (PIC) formation, initiation, pause-release, elongation and termination) (Fuda et al., 2009; Nagel and Taatjes, 2025) are regulated by enhancers to increase target gene expression remains unresolved. It has been proposed that, through the cooperative binding of TFs, enhancers recruit co-activators (e.g. MED, p300, Brd4) that are then “delivered” to the target promoters to predominantly control transcription initiation (Larke et al., 2021). However, these co-activators do not exclusively regulate PIC formation and initiation, but also other steps along the transcriptional cycle (Nagel and Taatjes, 2025). Accordingly, some enhancers seem to preferentially control RNA Pol II pause release and elongation rather than initiation (Henriques et al., 2018).

One of the reasons that could explain why, despite decades of extensive research, some fundamental aspects of enhancer function remain elusive is that enhancers represent a large and heterogeneous group of *cis*-regulatory elements. In this regard, previous studies indicate that there are multiple types of enhancers that differ in their underlying genetic architecture and regulatory properties (e.g. typical or “Core” enhancers, CpG-rich enhancers, Super-enhancers, Locus Control Regions, CTCF-associated enhancers, YY1-associated enhancers, LDB1-bound enhancers, Epromoters, *etc.*)(Aboreden et al., 2025; Boija et al., 2018; Crispatzu et al., 2021; Dao et al., 2017; Grosveld et al., 1987; Kubo et al., 2021; Pachano et al., 2021). Among them, CTCF-associated enhancers and Super-Enhancers (SE) are considered particularly important in the establishment of cell type-specific gene expression programs in the context of both development and human disease (Boija et al., 2018; Bower and Kvon, 2025; Kubo et al., 2021; Ren et al., 2017; Sabari et al., 2018):

i. CTCF-associated enhancers: a significant number of enhancer-promoter pairs exhibiting physical proximity are associated with nearby CTCF sites (CBS) (Bower and Kvon, 2025; Kubo et al., 2021). This suggests that, besides its well established role in insulation, CTCF might also work as a long-range facilitator that increases the contact frequency between enhancers and promoters (Rinzema et al., 2022). This facilitator role has been confirmed at several loci (e.g. *SHH, SOX9, PITX1, MYC*) containing ultra-long range enhancers (>400 Kb from target gene), where either the deletion or insertion of CBS regulates both enhancer-promoter physical proximity and gene expression (Bower and Kvon, 2025; Kragesteen et al., 2018; Lettice et al., 2002; Long et al., 2020; Yanchus et al., 2022). However, this facilitator role might not be universal, as the deletion of CBS associated with more proximal enhancers has a limited impact on target gene expression (Chakraborty et al., 2023; Ealo et al., 2024; Pachano et al., 2021) and the global depletion of CTCF does not have major effects on either enhancer-promoter contacts or gene expression programs (Nora et al., 2017).
ii. “Super-enhancers” (SEs): SE were originally defined as large clusters of nearby enhancers that recruit particularly high levels of co-activators, such as Mediator, and robustly control the expression of cell identity genes (Boija et al., 2018, 2018; Hnisz et al., 2013; Whyte et al., 2013). It has been suggested that SEs display emergent regulatory properties that go beyond the simple additive effect of their components (i.e. “core” enhancers). Furthermore, the strong regulatory capacity of SEs has been attributed to the establishment of low-affinity multivalent interactions between TFs and co-activators that are rich in intrinsically disorder regions (IDRs), which in turn can lead to the formation of transcriptional condensates through liquid-liquid phase separation. Current models propose that the formation of such condensates locally increases the concentration of the transcriptional machinery and ensure the robust expression of the SE target genes (Boija et al., 2018; Sabari et al., 2018). However, the emergent rather than additive properties of SE as well as the importance of condensates for their strong regulatory activity remain controversial (Chen et al., 2023; Meeussen et al., 2023; Shimizu et al., 2018; Trojanowski et al., 2022; Wu et al., 2022).

Previous studies in which different types of enhancers were investigated suffered from potential confounding factors (e.g. differences in model organism, cell type, developmental stage, TAD/regulatory domain, enhancer-promoter linear distance, type and/or epigenetic state of target promoter, *etc.*), making it difficult to discern the distinct regulatory properties and mechanisms of action of each enhancer type. To minimize this problem, here we used a synthetic biology approach to systematically compare the regulatory properties of major enhancer types (i.e. “typical” enhancer, CTCF-associated enhancer and SE/enhancer cluster) under identical conditions and using a wide variety of methods (i.e. ChIP-seq, Capture-C, single-molecule RNA-FISH, IF+RNA-FISH). Chiefly, our results argue against a general role for CTCF as a facilitator of long-range enhancer-gene communication, while providing novel insights into the mechanisms whereby enhancer clustering increases target gene expression. Namely, we propose that the emergent regulatory properties of enhancer clusters (e.g. SEs) can not be simply explained by changes in enhancer-promoter contact frequency or the formation of transcriptional condensates, but rather entail a strong increase in the fraction enhancer-promoter contacts that are transcriptionally productive.

## Results

### Synthetic engineering of different enhancer types with a shared “core” enhancer

We recently implemented a genetic engineering approach to introduce different types of synthetic enhancers at a given locus and compare their regulatory properties under highly similar conditions (Pachano et al., 2021). Briefly, an evolutionary conserved enhancer located 35 Kb downstream of *Sox1* (*Enh Sox1(+35))* that becomes active (i.e. gains p300 and H3K27ac) and controls *Sox1* induction upon differentiation of mESC into neural progenitor cells (NPC) was used as a “core” enhancer from which additional enhancer types were built (Fig1A-C) (Cruz-Molina et al., 2017; Pachano et al., 2021). Then, the different synthetic enhancers were inserted in mESC at the same distance from a developmental gene (i.e.100 Kb downstream of *Gata6*) with a promoter responsive to distal enhancers (Pachano et al., 2021) (Fig 1D). By inserting the synthetic enhancers within the regulatory domain of a gene that is inactive in mESCs and NPCs (i.e. *Gata6*) (Cruz-Molina et al., 2017) (Fig 1C), and, thus, devoid of active enhancers in these cell types (Fig S1A), changes in target gene expression that occur upon NPC differentiation can be attributed solely to the inserted enhancer sequences. In principle, this reductionist approach should enable a more accurate comparison between different enhancer types by minimizing potential confounding factors (e.g. model organism, cell type, developmental stage, TAD/regulatory domain, enhancer-promoter linear distance, type and/or epigenetic state of target promoter, analytical methods, *etc.*) that exist among previous studies in which different types of enhancers were investigated.

**Fig 1:**
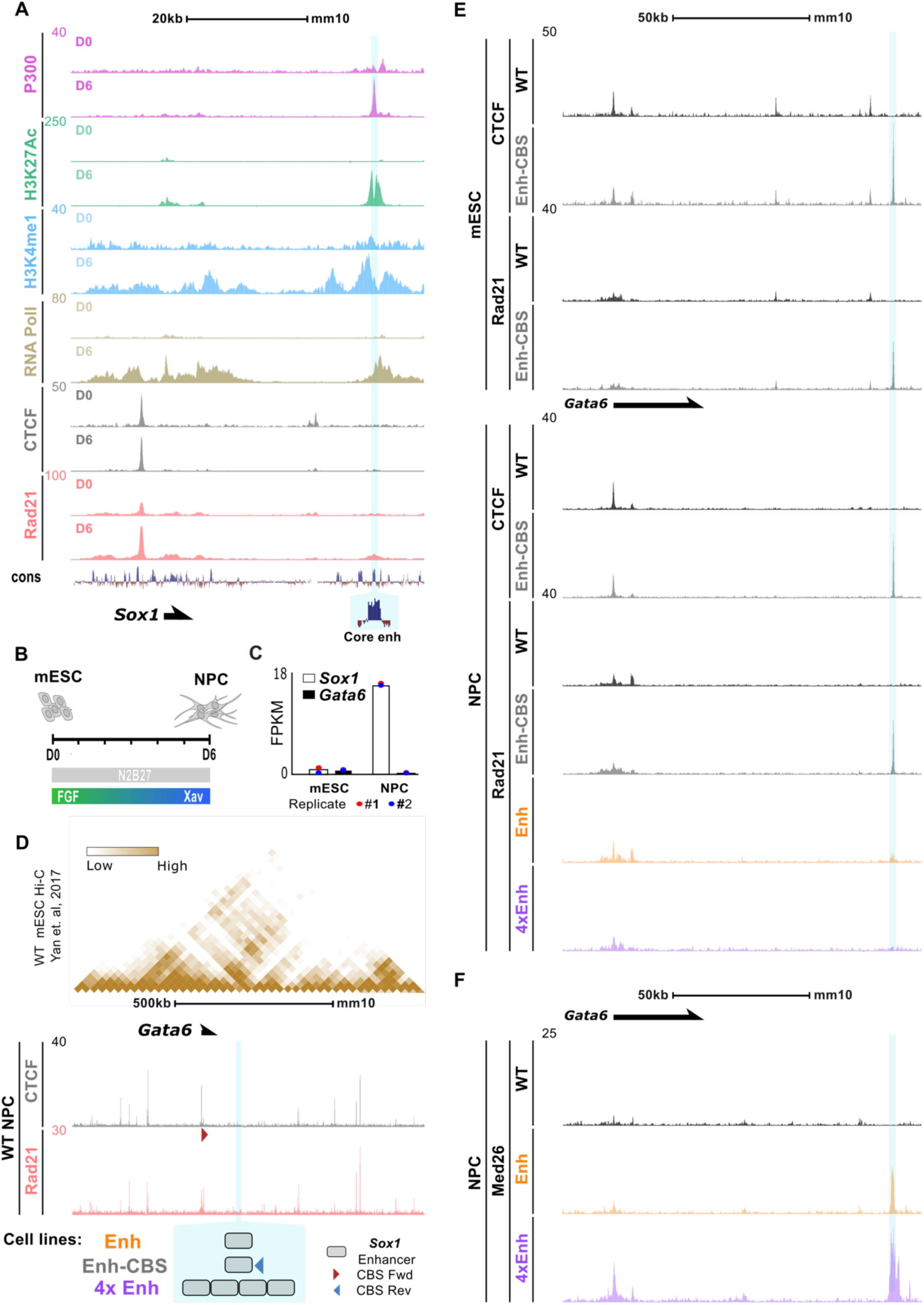
Synthetic enhancers exhibit molecular properties that reflect their distinct genetic architectures. **(A)** Genome browser tracks displaying ChIP-seq signals for the indicated coactivators, histone marks, CTCF and RAD21 in mESCs and Neural Progenitor Cells (NPCs) at the Sox1 locus. Vertebrate conservation is shown at the bottom (cons). The position of the Sox1(+35) enhancer used as the “core” element to build different types of synthetic enhancers is shaded in blue. **(B)** Illustration depicting the in vitro differentiation of mESC into NPC. **(C)** Expression levels (FPKM) of Gata6 and Sox1 in mESC and NPC (Cruz-Molina et al., 2017). **(D)** mESC Hi-C data at the Gata6 locus (Yan et al., 2018) on top. Below, ChIP-seq tracks for CTCF and Rad21 are shown. The blue highlighted region indicates the insertion site of the synthetic enhancers depicted underneath. **(E)** CTCF and Rad21 ChIP-seq signals at the Gata6 locus are shown in WT mESC and CBS-Enh mESC (top) as well as upon their differentiation into NPC (bottom). Rad21 ChIP-seq profiles in NPC are also shown for the Enh and 4xEnh cell lines (bottom). CTCF and Rad21 ChIP-seq profiles in WT and Enh- CBS cells are presented as the average from two independent replicates (N = 2), while the Rad21 ChIP-seq profiles in the Enh and the 4xEnh cells are based on a single replicate (N = 1). The insertion site of the synthetic enhancers is shaded in blue. **(F)** Average MED26 ChIP-seq signals (two biological replicates per cell line) at the Gata6 locus are shown for WT, Enh and 4xENH NPC. Alignment of the ChIP-seq data was performed using mm10-based custom reference genomes including the inserted enhancers for each of the different cell lines.

Here we used the synthetic engineering approach described above to gain insights into the regulatory mechanisms employed by three enhancer types (i.e. “typical” or “core” enhancers, CTCF-associated enhancers, and SEs/enhancer clusters) and clarify ongoing debates regarding: (i) the role of CTCF as facilitator of enhancer-promoter communication and (ii) the emergent properties attributed to SEs. The *Enh Sox1(+35)* described above was used as a “core” enhancer to design (Fig 1D): (i) Enh: a typical enhancer containing the “core” enhancer alone (Pachano et al., 2021), (ii) Enh-CBS: a CTCF-associated enhancer containing a CBS downstream of the “core“ enhancer and (iii) 4xEnh: an enhancer cluster resembling a SE and consisting of four tandem repeats of the “core” enhancer. The Enh-CBS was generated by adding a CBS in reverse orientation immediately downstream of the “core“ enhancer. A CBS located upstream of the *Shh* gene was selected based on its capacity to stall cohesin both in mESC and NPC (Fig S1B). On the other hand, the decision to build an enhancer cluster with four tandem copies of the “core” enhancer (i.e. 4xEnh) that could somehow mimic the regulatory properties of SEs was based on the previously described genetic features of this enhancer type: (i) SEs whose individual components have been genetically dissected consist of three to five regulatory elements (e.g. α-globin LCR/SE: five; *Sox2* SE: four; *Prdm14* SE: five; *Pou5f1* SE: three; *WAP*: three); (ii) SEs globally identified in various cell types have, on average, around four individual components per SE (e.g. hESC: 3.31 elements; hMSC: 4.31 elements; SEdb2.0 (Jiang et al., 2019)). Next, we generated transgenic mESC lines in which the previous synthetic enhancers (*i.e.* Enh (already generated and characterized in Pachano et al., 2021), Enh-CBS and 4xEnh) were inserted 100 Kb downstream of the *Gata6* TSS (Fig 1D; Fig S1C-F), a linear distance at which “typical” enhancers need assistance from nearby tethering elements and/or the cohesin complex to robustly drive target gene expression (Pachano et al., 2021; Rinzema et al., 2022). Importantly, the *Gata6* promoter contains a CBS with a convergent orientation (i.e. forward; red triangle in Fig 1D) with respect to the Enh- CBS (i.e. reverse), which in principle should facilitate the establishment of a loop between the *Gata6* promoter and the Enh-CBS (de Wit et al., 2015; Fudenberg et al., 2016; Nora et al., 2017; Rao et al., 2017; Sanborn et al., 2015; Schwarzer et al., 2017).

### Initial validation of the key molecular properties of the Enh-CBS and 4xEnh synthetic enhancers

To validate whether the Enh-CBS and 4xEnh displayed the expected molecular features when inserted within the *Gata6*-TAD, ChIP-seq experiments against CTCF, RAD21 (a subunit of the Cohesin complex) and MED26 (a subunit of the Mediator complex) were performed in the transgenic mESC lines described above (Fig 1E-F). The addition of the CBS to the “core” enhancer resulted in strong binding of CTCF in both mESC and NPC (Fig 1E). Accordingly, the Enh-CBS showed significantly increased stalling of cohesin (i.e. RAD21 ChIP-seq signal) relative to the Enh alone or the 4xEnh in NPC (Fig 1E). On the other hand, increased recruitment of Mediator to the 4xEnh cluster was confirmed by comparing MED26 ChIP-seq signals in NPC derived from the *4xEnh* and *Enh* mESC lines (Fig 1F), which revealed that the 4xEnh recruited ∼3 times more MED26 than the Enh alone.

### Enhancer clustering leads to a synergistic increase in long-range regulatory activity, while the addition of the CBS does not facilitate enhancer function

To determine the regulatory activity of each enhancer type, we measured *Gata6* expression by RT-qPCR upon differentiation of the transgenic mESC lines into NPC. Expression analyses for the *Enh-CBS* and *4xEnh* lines were performed using at least three independent clonal mESC lines for each genotype, while for the previously characterized *Enh* cells (Pachano et al., 2021) a single clonal mESC line was used. Surprisingly, *Gata6* expression was very similar in *Enh* and *Enh-CBS* NPC (*Enh:* 14.6 ± 4.2 fold-change over WT, p-value = 3.6e-5; *Enh-CBS*: 14.3 ± 4.5 fold-change over WT, p-value = 3.5e-5) (Fig 2A), thus indicating that the binding of CTCF to the “core” enhancer had no effect on *Gata6* expression (p-value > 0.05). In stark contrast, *Gata6* expression in cells containing the 4xEnh was almost 180 times higher than in WT NPC (*4xEnh*: 179 ± 46 fold-change over WT, p-value = 2.4e-9) (Fig 2A). Therefore, despite only having four copies of the “core“ enhancer, the 4xEnh cluster showed a synergistic 13-fold increase in *Gata6* expression relative to the Enh alone (p-value = 4.06e-7). This further supports that our synthetic enhancer cluster displays emergent regulatory properties similar to those previously proposed for SEs (Boija et al., 2018; Sabari et al., 2018).

**Fig 2:**
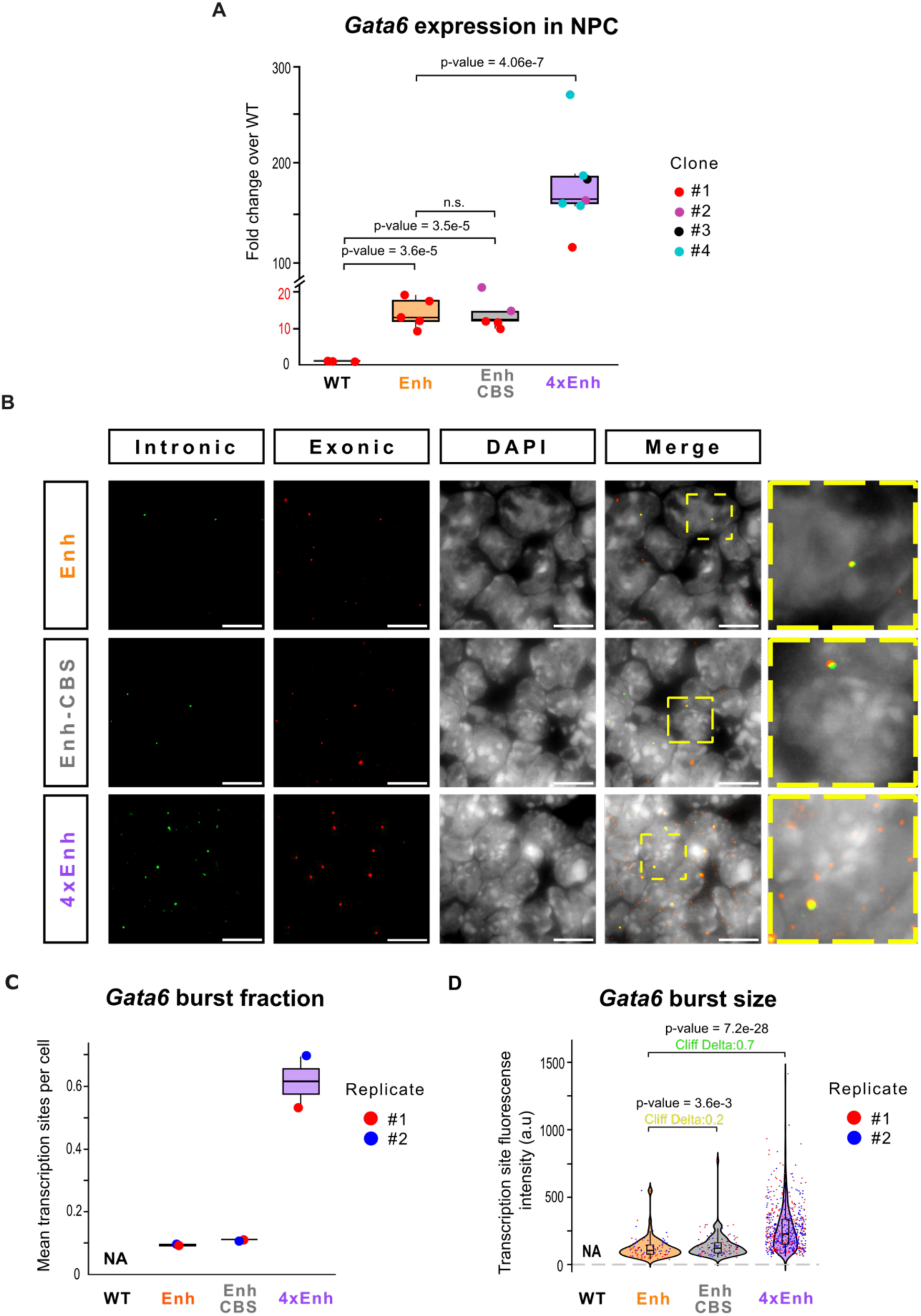
The 4xEnh cluster synergistically increases Gata6 expression and burst fraction, while the CBS has a mild effect on the regulatory properties of the “core” enhancer. **(A)** Boxplots showing Gata6 expression levels as measured by RT-qPCR in WT NPC and NPC derived from the transgenic mESC lines with the synthetic enhancers. Expression values for each transgenic cell line are normalized with respect to the WT cells using two housekeeping genes (Hprt and Eef1a) as loading controls. Biological replicates are represented by dots, while the colors indicate independent clonal cell lines. P-values were calculated using two- sided unpaired t-tests (p > 0.05: NS (not significant). **(B)** Representative images of single- molecule RNA FISH experiments for Gata6 in the indicated cell lines. Intronic (green) and exonic (red) signals, along with DAPI staining, are shown to visualize nascent mRNA foci. For visualization purposes, five Z-stacks (0.5 μm per stack) were merged, and intron and exon channels were adjusted to equal intensity levels. A merged image of all three channels is included to highlight the nuclear overlap between intronic and exonic signals. Zoom-in views of the regions highlighted in yellow are shown to the far right. Scale bar: 10 μm. **(C)** Boxplot depicting the mean number of Gata6 nascent transcription sites (i.e. foci in which the intronic and exonic signals overlap) per cell (i.e. burst fraction) for the indicated cell lines upon analysis of 1000 cells per genotype across two independent replicates. **(D)** Violin plot showing the distribution of intronic signal intensities (i.e. burst size) at Gata6 nascent transcription sites for the indicated cell lines. Each dot represents the intronic signal intensity of a single nascent transcription site. Data were collected from 1000 cells per genotype across two independent replicates. P-values were calculated using unpaired two-sided Wilcoxon tests with Holm correction. Effect size was calculated using the Cliff’s delta statistic (green: large effect size; yellow: small effect size).

### Enhancer clustering synergistically increases target gene expression by preferentially regulating burst fraction and, to a lesser extent, burst size

The synergistic effect in transcriptional output observed in cells containing the 4xEnh cluster prompted us to characterize transcriptional bursting parameters associated to each enhancer type. To do so, we used single-molecule RNA fluorescence in situ hybridization (smRNA-FISH), a method that enables the measurement of burst fraction and burst size in fixed cells (Chubb et al., 2006; Femino et al., 1998; Raj et al., 2006). Burst fraction, which is related to and used as a measure of burst frequency, refers to the number of alleles transcribing per cell at a given time, while burst size corresponds to the number of RNA molecules produced per burst (Bartman et al., 2016; Raj et al., 2006).

smRNA-FISH experiments were performed using both intronic and exonic probes targeting *Gata6* in NPC carrying the different synthetic enhancers (Fig 2B). For each enhancer type, a representative clonal ESC line was used in two independent biological replicates, resulting in the analysis of approximately 1000 cells per genotype. Similarly to the previous RT-qPCR experiments, the addition of the CBS to the “core” enhancer had a very mild effect on either burst fraction (*Enh:* 0.09 ± 0.005; *Enh-CBS*: 0.11 ± 0.003) (Fig 2C) or burst size (*Enh:* 117 ± 76; *Enh-CBS*: 138 ± 86) (Fig 2D) compared to the “core” enhancer alone. In contrast, the 4xEnh cluster showed a ∼6-fold increase in burst fraction (*4xEnh*: 0.610 ± 0.11) and a ∼2-fold increase in burst size (*4xEnh*=260 ± 157) relative to the “core” enhancer alone. Overall, these results indicate that the 4xEnh cluster synergistically increases *Gata6* expression by preferentially increasing the burst fraction and, to a lesser, but not negligible extent, also the burst size (Fig 2D; *4xEnh* vs *Enh*: p-value = 7.2e-28, Cliff Delta=0.7). Therefore, although enhancers are typically considered to only control burst fraction/frequency (Bartman et al., 2016; Fukaya et al., 2016; Larson et al., 2013; Larsson et al., 2019; Walters et al., 1995), our findings suggest that certain enhancers (e.g. enhancer clusters/SEs) might also modulate the burst size.

### Enhancer-associated CTCF sites increase intra-TAD insulation while having a mild effect on enhancer-promoter contact frequency

Intrigued by the lack of long-range regulatory differences between the Enh-CBS and the Enh, we next performed Capture-C experiments to evaluate whether, despite not having a major effect on *Gata6* expression, the binding of CTCF next to the “core” enhancer could still affect the 3D chromatin organization of the *Gata6*-TAD. Capture-C experiments were performed in NPC using as viewpoints either the *Gata6* promoter (Fig 3A) or the enhancer insertion site located 100kb downstream of the promoter (Fig 3B). Regardless of the considered viewpoint, the interaction frequency between the *Gata6* promoter and the insertion site exhibited a ∼3.5-fold increase in *Enh* cells relative to the WT control (Fig 3A-B). Importantly, such interactions were either unchanged (*Gata6* promoter viewpoint; Fig 3A), or slightly higher (∼1.4-fold; insertion site viewpoint; Fig 3B) in *Enh-CBS* compared to *Enh* cells, thus in agreement with the lack of significant differences in *Gata6* expression levels between both cell lines (Fig 2A). Nevertheless, the addition of the CBS to the “core” enhancer did have a significant impact on the 3D chromatin organization of the *Gata6*-TAD that could be best appreciated upon subtraction of the WT Capture-C signals from the signals obtained in the transgenic cell lines (i.e. subtraction tracks in Fig 3A-B). Namely, the addition of the CBS created a new subdomain within the *Gata6*-TAD spanning from the insertion site up to the 5′ TAD boundary located upstream of *Gata6*. As a result, in *Enh-CBS* cells the *Gata6* promoter and the inserted enhancer showed increased interaction frequencies with regions located within the new subdomain, including clear interactions with CTCF peaks demarcating the 5’ boundary of the *Gata6*-TAD. In contrast, interactions with regions located beyond (i.e. 3’) the insertion site were reduced.

**Fig 3:**
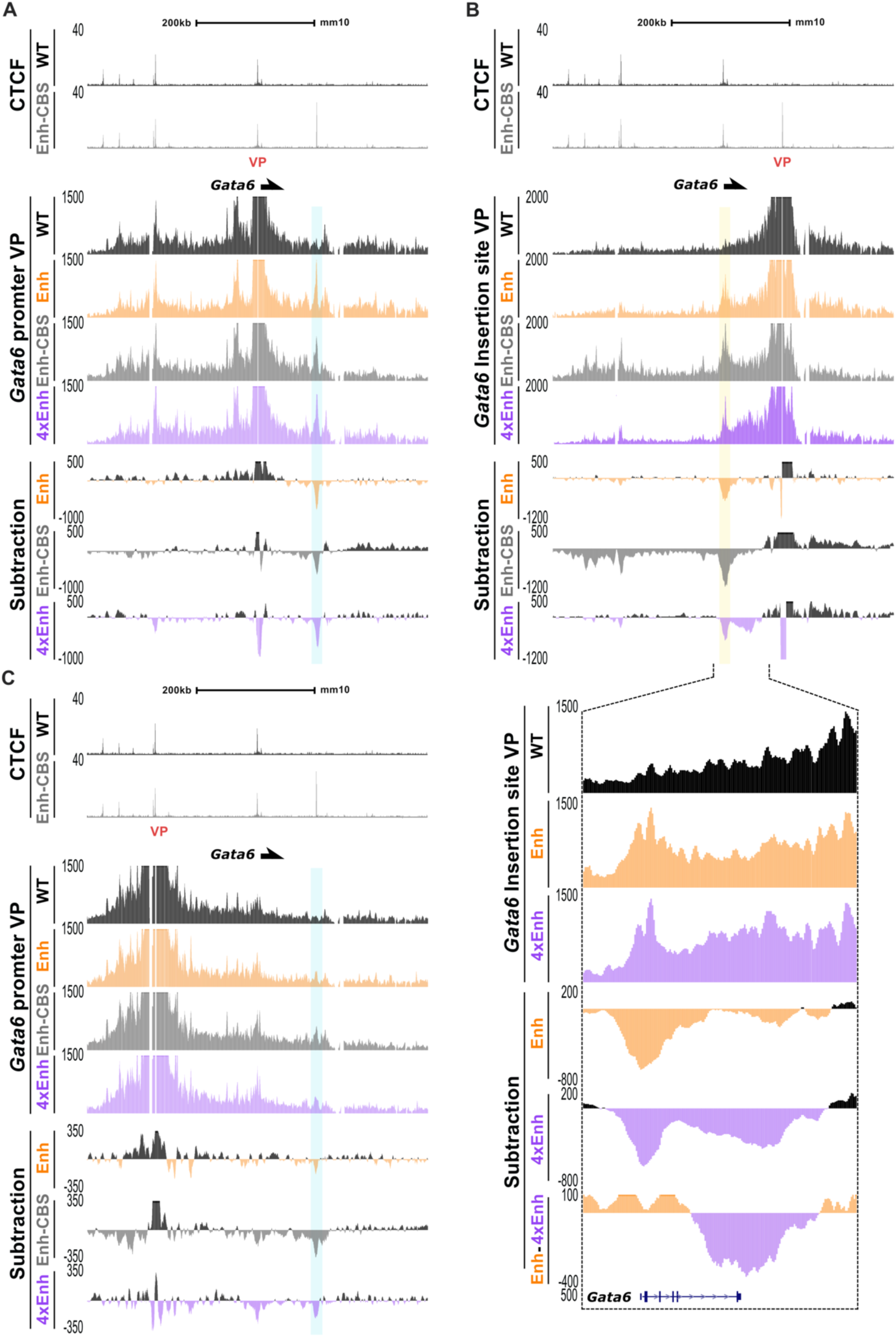
Capture-C experiments reveal that the three synthetic enhancers display similar contact frequencies with the Gata6 promoter. **(A)** The upper tracks display CTCF ChIP- seq profiles at the Gata6 locus in WT and Enh-CBS NPC. Below, Capture-C profiles using the Gata6 promoter as viewpoint (VP) are shown in NPC derived from the indicated cell lines. For each cell line, the Capture-C profiles were generated by averaging the signals from two independent replicates (N = 2). Underneath, subtraction tracks are presented in order to highlight the differences in Capture-C signals between each synthetic enhancer cell line and the WT cells around the Gata6 locus. **(B)** Same as in (A) but using the synthetic enhancer insertion site as a viewpoint (VP). The tracks at the bottom show a zoomed-in view of the Capture-C signals around the Gata6 gene body to better visualize the contacts between the 4xEnh cluster and the gene body. **(C)** Same as in (A) but using one of the CTCF peaks located within the 5′boundary of the Gata6-TAD as viewpoint. In (A-C), the insertion site is highlighted in blue and the Gata6 promoter in yellow.

The contacts between the inserted Enh-CBS, the *Gata6* promoter and the CTCF peaks overlapping the 5’ TAD boundary that are preferentially observed in *Enh- CBS* cells suggest that the sub-domain that emerges upon addition of the CBS might display a hub-like architecture with nested multi-way interactions (Karpinska et al., 2025). To further test this possibility, additional Capture-C experiments were performed using as a viewpoint one of the prominent CTCF peaks overlapping the 5′ TAD boundary of the *Gata6*-TAD. Interestingly, in *Enh-CBS* cells, the 5’ TAD boundary showed higher contact frequency with both the *Gata6* promoter and the insertion site in comparison to either WT or *Enh* cells (Fig 3C). Collectively, our results suggest that the addition of the CBS to the “core” enhancer generates an insulated sub-domain with a hub-like architecture spanning from the insertion site to the 5′ boundary of the *Gata6*-TAD, which, nevertheless, has a rather modest effect on either enhancer-*Gata6* promoter contact frequency or *Gata6* expression (Fig 2). Altogether, our data supports a role for CTCF in insulation and chromatin hub formation, rather than as a general facilitator of enhancer-promoter communication and enhancer function.

### The synergistic effect of the 4xEnh cluster on target gene expression does not involve changes in enhancer-promoter contact frequency

In contrast to the Enh-CBS, the 4xEnh cluster synergistically increased *Gata6* expression by preferentially boosting the fraction of cells expressing *Gata6*. Therefore, we wondered whether, as previously proposed for other enhancers, the 4xEnh cluster controlled the *Gata6* burst fraction by contacting more frequently with its promoter (Bartman et al., 2016; Tünnermann et al., 2025; Zuin et al., 2022). To test this, Capture-C experiments similar to the ones described in the previous section were also performed in *4xEnh* NPC. Remarkably, using either the *Gata6* promoter or the insertion site as viewpoints revealed that the contact frequency between the 4xEnh and the *Gata6* promoter was similar to the one observed for the “core” enhancer alone (Promoter viewpoint: *Enh*: 3.48 and *4xEnh*: 4.02; Insertion site viewpoint: *Enh*: 3.32 and *4xEnh*: 2.9; fold increase relative to WT cells) (Fig 3A-B). Furthermore, in addition to the discrete contact with the *Gata6* promoter, the 4xEnh showed increased interactions over the whole *Gata6* gene body in comparison to both *Enh* and WT cells (Fig 3B). Similarly, interactions between the *Gata6* promoter and the *Gata6* gene body were also stronger in *4xEnh* cells when using the *Gata6* promoter as a viewpoint (Fig S2A). The contacts between the 4xEnh cluster, the *Gata6* promoter and its gene body are in agreement with previous reports suggesting that, as genes undergo transcription elongation, enhancers and their target promoters establish dynamic interactions throughout the gene bodies that track with the elongating RNA Pol II (Lee et al., 2015). Taken together, our results suggest that the 4xEnh cluster does not increase *Gata6* expression and burst fraction by increasing enhancer-promoter contact frequency but may instead do so by increasing the proportion of productive contacts leading to transcription elongation.

### Enhancer clustering increases coactivator recruitment in a sub-additive manner despite its synergistic effect on transcriptional output

SEs consist of clusters of enhancers that are densely occupied by TFs, coactivators and H3K27ac (Hnisz et al., 2013; Whyte et al., 2013). To further characterize the potential mechanisms behind the emergent regulatory properties of the 4xEnh cluster, we generated ChIP-seq profiles in WT, *Enh* and *4xEnh* NPCs for key histone marks (i.e. H3K4me2, H3K4me3, H3K27ac and H3K27me3)(Heintzman et al., 2009) and co-activators (MED26, BRD4, p300)(Hnisz et al., 2013; Visel et al., 2009; Whyte et al., 2013) previously associated with transcriptional regulation and enhancer function. As the addition of the CBS to the *“core”* enhancer did not have any major effect on *Gata6* expression, ChIP-seq experiments were not performed in *Enh- CBS* cells.

Overall, the ChIP-seq profiles for the histone marks around the *Gata6* promoter were consistent with the *Gata6* expression levels measured in the different cell lines. Namely, the repressive H3K27me3 mark was slightly decreased at the *Gata6* promoter in *Enh* cells and further diminished in *4xEnh* cells in comparison to the WT control. In contrast, for the active histone modifications, the ChIP-seq data showed that H3K27ac (*Enh*: 2 ± 0.44 fold-change over WT; *4xEnh*: 12.3 ± 0.96 fold change over WT) and H3K4me3 (*Enh*: 2.55 ± 0.08 fold-change over WT; *4xEnh*: 7.51 ± 2.48 fold-change over WT) levels around the *Gata6* promoter were significantly higher in *4xEnh* cells than in both *Enh* and WT cells, while for H3K4me2 a modest ∼2-fold increase over WT was observed in both *Enh* and *4xEnh* cells (Fig 4A-B). Interestingly, H3K4me3 levels in *4xEnh* cells were particularly increased downstream of the *Gata6* TSS, which given the proposed role for this histone mark in RNA Pol II pause release, further suggests that the 4xEnh cluster preferentially increases productive transcription and elongation (Wang et al., 2023). Regarding the histone modification levels around the enhancer insertion site, while both *Enh* and *4xEnh* cells exhibited a ∼20 fold increase in H3K4me2 relative to the WT, the levels of H3K27ac were higher in *4xEnh* cells (*Enh*: 14.55 ± 2.5 fold-change over WT; *4xEnh*: 41.19 ± 21.19 fold-change over WT) (Fig 4A-B), thus further supporting that our synthetic enhancer cluster recapitulates the main molecular features of SEs (i.e. high levels of Mediator (Fig 1F) and H3K27ac (Fig 4A-B) (Hnisz et al., 2013; Whyte et al., 2013).

**Fig 4:**
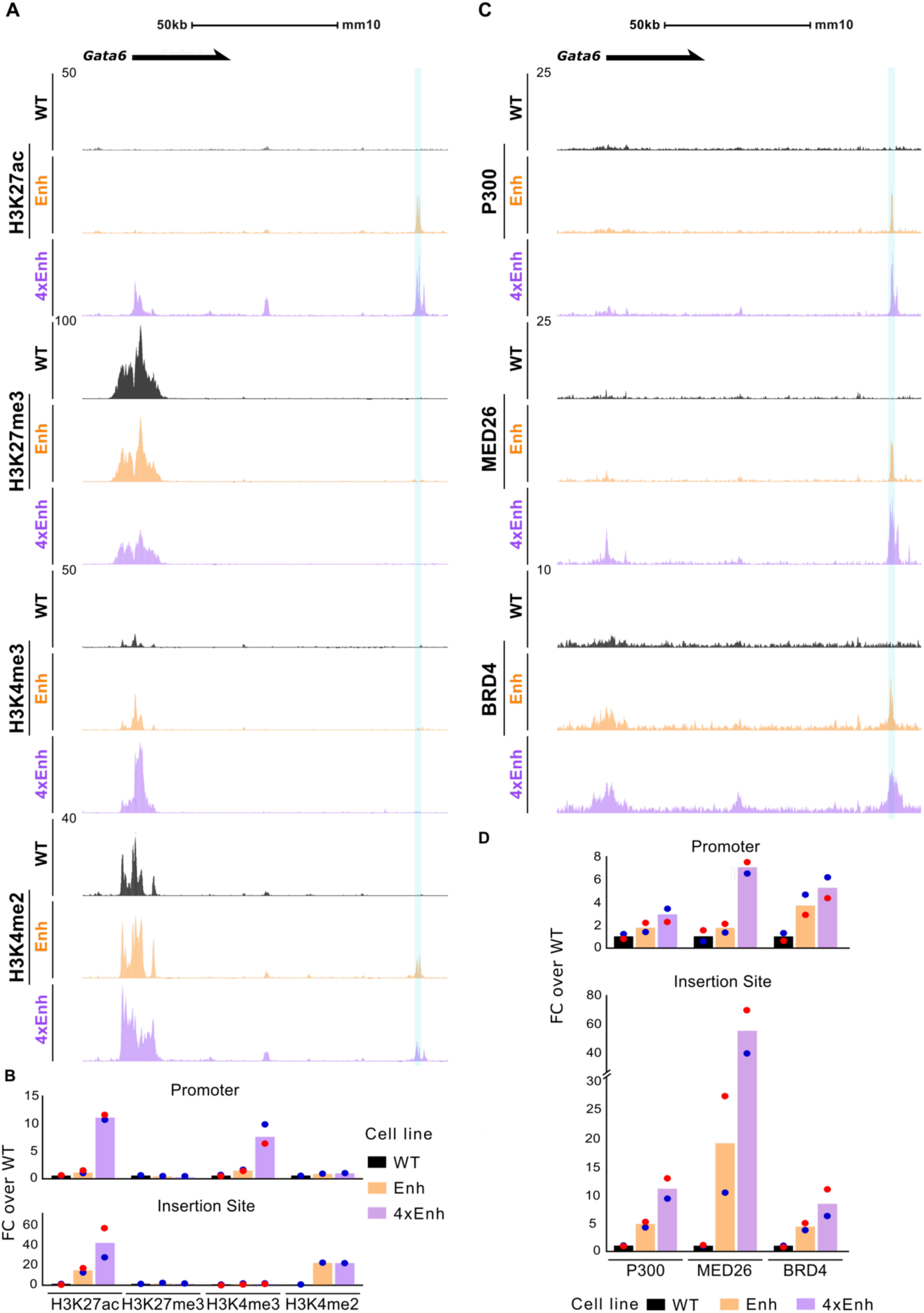
Sub-additive co-activator recruitment at the 4xEnh cluster. **(A)** Genome browser tracks showing ChIP-seq profiles for the indicated histone marks at the Gata6 locus in WT NPC and NPC with the different synthetic enhancers. H3K27ac and H3K4me3 tracks represent the average signal from two independent replicates (N = 2), while H3K27me3 and H3K4me2 are based on a single replicate (N = 1). **(B)** Quantification of the ChIP-seq signals for the histone marks shown in (A) at either the Gata6 promoter (top) or the synthetic enhancer insertion site (bottom). For the Gata6 promoter, a region spanning ± 2.5 kb around the TSS was used. For the insertion site, a window extending 500 bp upstream and 2.5 kb downstream of the insertion point was considered (Supplementary Data 1). **(C)** Genome browser tracks showing the ChIP-seq profiles for the indicated co-activators (i.e. p300, MED26 and BRD4) at the Gata6 locus in WT NPC and NPC with the different synthetic enhancers. The ChIP-seq tracks represent the average signals from two independent replicates (N=2) for each co- activator and cell line. **(D)** Quantification of the ChIP-seq signals for the co-activators shown in (C) at either the Gata6 promoter (top) or the synthetic enhancer insertion site (bottom). For the Gata6 promoter, a region spanning +/-300bp around the TSS was used. For the insertion site, a window extending 500 bp upstream and 2.5 kb downstream of the insertion point was considered (Supplementary Data 1). In (B) and (D), the quantifications of the ChIP-seq signals in the synthetic enhancer cell lines are presented as fold-changes relative to the ChIP-seq signals in WT cells.

As described in previous sections, (Fig 1F), the 4xEnh cluster recruited ∼3 to 4 times more MED26 than the “core” enhancer alone at the insertion site and the TSS, respectively (Fig 4C-D) (TSS - *Enh*: 1.79 ± 0.56 fold-change over WT; *4xEnh*: 7.05 ± 0.72 fold-change over WT; Insertion site - *Enh*: 19.2 ± 11.88 fold-change over WT; *4xEnh*: 55.18 ± 20.29 fold-change over WT). Next, we investigated whether the 4xEnh cluster also recruited higher levels of other coactivators (i.e. p300, BRD4) that, although not as pronouncedly as for Mediator, are also enriched at SEs compared to typical enhancers (Boija et al., 2018; Hnisz et al., 2013; Sabari et al., 2018). Both p300 (*Enh*: 1.7 ± 0.61 fold-change over WT; *4xEnh*: 2.9 ± 0.86 fold-change over WT) and BRD4 (*Enh*: 3.8 ± 1.3 fold-change over WT; *4xEnh*: 5.29 ± 1.2 fold-change over WT), exhibited a mild increase around the *Gata6* promoter in both *Enh* and *4xEnh* cells compared to their WT counterparts (Fig 4C-D). Furthermore, at the insertion site, the *4xEnh* cells showed a more prominent (∼2 fold) increase than the *Enh* cells for both coactivators (P300 - *Enh*: 5.43 ± 0.76 fold-change over WT; *4xEnh*: 11.45 ± 2.43 fold- change over WT; BRD4 - *Enh*: 4.4 ± 0.92 fold-change over WT; *4xEnh*: 8.57 ± 3.3 fold- change over WT) (Fig 4C-D).

Taken together, these ChIP-seq experiments revealed that, in comparison to the “core” enhancer alone, the 4xEnh cluster recruits high levels of multiple co- activators (i.e. Mediator, p300, BRD4), which, nevertheless, increase in a sub-additive manner in comparison to the “core” enhancer alone (i.e. 4xEnh vs Enh fold-changes < 4). This suggests that the strong regulatory activity and synergistic transcriptional output of the 4xEnh cluster might arise through the sub-additive recruitment of multiple coactivators, which together could bypass some of the rate limiting steps of transcription (Fuda et al., 2009; Wissink et al., 2019).

### The strong regulatory activity of the cluster weakly correlates with the formation of transcriptional condensates

The emergent regulatory properties of SEs have been attributed to the strong recruitment of co-activators rich in IDRs, such as Mediator or BRD4, which can facilitate the formation of condensates that enable the robust expression of major cell identity genes (Boija et al., 2018; Hnisz et al., 2013; Sabari et al., 2018; Whyte et al., 2013). Due to the large size of these transcriptional condensates (Du et al., 2024), enhancers and promoters might co-exist within them without establishing close contacts as measured by 3C-related methods (Yang and Hansen, 2024), thus potentially explaining how the 4xEnh cluster might increase *Gata6* expression without increasing the contact frequency with its promoter (Fig 2A and 3A-B). To test this possibility, we next compared the ability of the “core” enhancer and the 4xEnh cluster to form transcriptional condensates by using a previously described strategy (Boija et al., 2018; Sabari et al., 2018) in which BRD4 immunofluorescence (IF) and *Gata6* smRNA-FISH assays are combined (Fig 5A). In agreement with previous reports (Boija et al., 2018; Sabari et al., 2018), we found that BRD4 condensates overlapped with *Gata6* nascent RNA-FISH foci more frequently in *4xEnh* cells than in *Enh* cells. Namely, cumulative analysis of BRD4 IF signals centered around the *Gata6* RNA-FISH foci in two independent replicates (n=100 *Gata6* nascent RNA foci in each cell line), showed that BRD4 was more enriched near the center of the RNA-FISH foci in *4xEnh* cells than in *Enh* cells (Fig 5B) as well as compared to randomly selected regions (see Methods for details) (Fig S3A). However, although quantification of the BRD4 IF signals within individual *Gata6* nascent RNA foci revealed that BRD4 signals were significantly higher in *4xEnh* cells than in *Enh* cells (p-value = 0.0017), the differences were subtle and strong *Gata6* RNA foci with low BRD4 signals in *4xEnh* cells were also observed (i.e. not overlapping a BRD4 condensate) (Fig S3B). Importantly, a more quantitative comparison between the nascent RNA-FISH and BRD4 IF signals at all the considered *Gata6* RNA-FISH foci revealed a weak positive correlation between both signals in both *Enh* and *4xEnh* cells (*Enh*: Spearman_rho = 0.06, p-value = 0.548; *4xEnh*: Spearman_rho = 0.137, p-value = 0.175) (Fig 5D).

**Fig 5:**
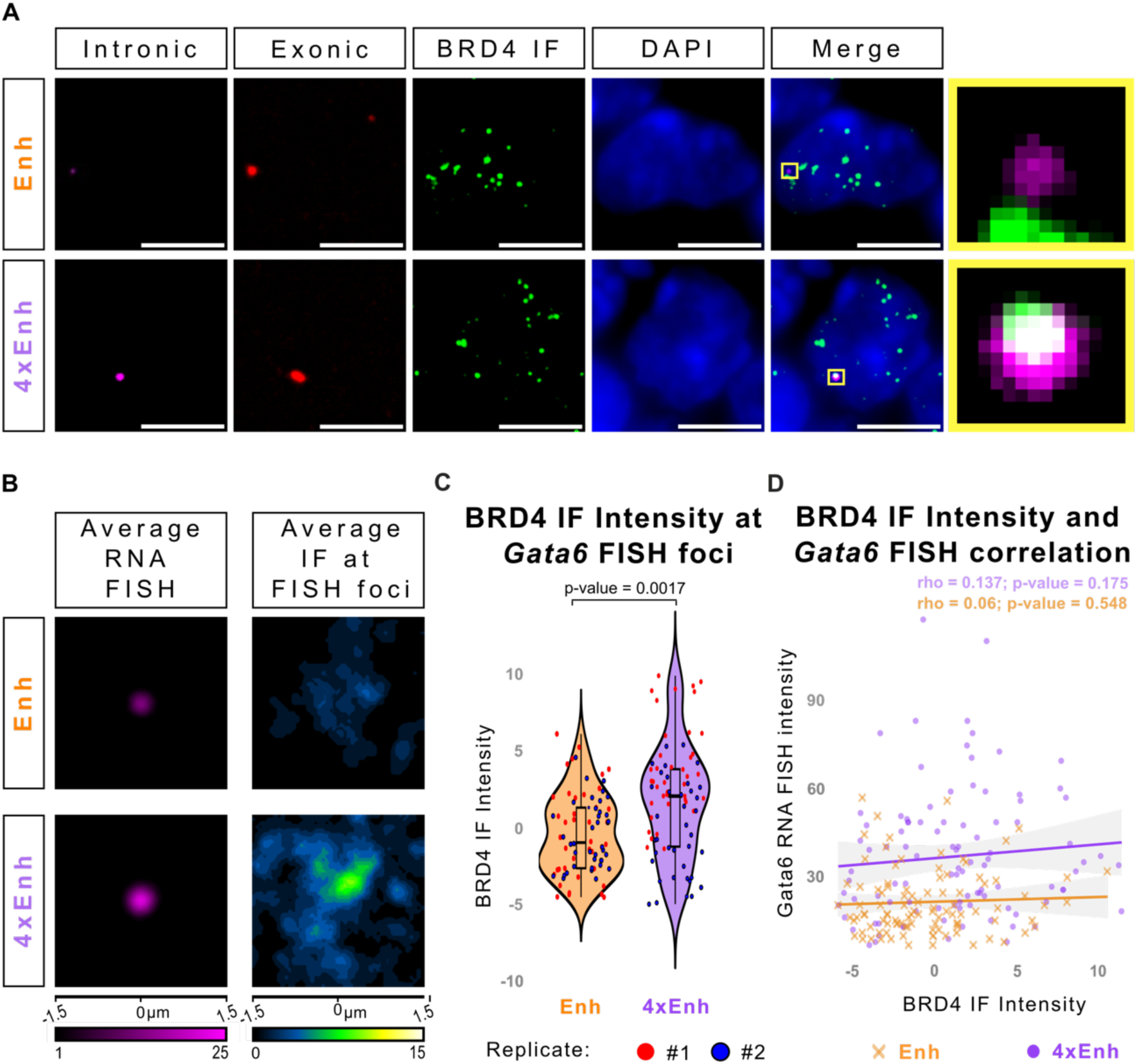
The emergent regulatory properties of the 4xEnh cluster are not explained by the formation of transcriptional condensates. **(A)** Co-localization of BRD4 immunofluorescence (IF) and Gata6 smRNA-FISH signals within selected cells. Individual panels display the BRD4 signal (green) and single-molecule RNA FISH signals targeting either intronic (magenta) or exonic (red) Gata6 regions in NPC derived from the indicated ESC lines. A composite image (Merge) including DAPI staining is shown to highlight the co-localization of the different signals. A zoomed-in view of the area highlighted by the yellow box is provided for clearer visualization of the overlap between IF and intronic signals. Scale bar: 5 µm. **(B)** Average intronic Gata6 smRNA-FISH signals (left) and BRD4 signals (right) centered at 100 Gata6 nascent transcription sites (i.e. foci in which the intronic and exonic signals overlap) identified in either Enh or 4xEnh NPC using two independent replicates per genotype. **(C)** Violin plots depicting the BRD4 fluorescence intensity at the Gata6 nascent transcription sites analyzed in B, after subtraction of the average background signal obtained at randomly selected regions (Fig S3A). P-values were calculated using unpaired two-sided Wilcoxon tests with Holm correction. **(D)** Correlation analysis between the intensities of the BRD4 signals and the intronic Gata6 smRNA-FISH signals at the Gata6 nascent transcription sites analyzed in (B) for either Enh or 4xEnh NPC.

The transcriptional condensate model previously proposed for SEs is largely based on correlative observations obtained upon comparison of a few genes that were regulated by either traditional enhancers or SEs and that, consequently, did not address whether condensates are causally involved in the robust expression of the SE target genes(Boija et al., 2018; Sabari et al., 2018). Overall, our synthetic engineering approach comparing the regulatory properties of the “core” enhancer and the 4xEnh cluster within the same locus indicates that, although the 4xEnh cluster shows a higher propensity to form condensates than the “core” enhancer, this does not explain its strong regulatory activity and synergistic effects on *Gata6* expression. Our findings are in agreement with recent reports questioning the importance of condensates for high transcriptional activity (Chen et al., 2023; Meeussen et al., 2023; Shimizu et al., 2018; Trojanowski et al., 2022; Wu et al., 2022).

### Enhancer clustering promotes RNA pol II release into productive elongation

In addition to facilitating the formation of condensates through LLPS, the co- activators showing increased recruitment to the 4xEnh cluster (i.e. p300, Mediator and BRD4) have been previously implicated in the control of different rate-limiting steps along the RNA Pol II transcription cycle, including PIC formation/initiation and RNA Pol II pause release (Altendorfer et al., 2022; Cramer, 2019; Narita et al., 2021). Therefore, we then assessed whether the 4xEnh cluster preferentially controlled any of those steps in comparison to the “core” enhancer alone by performing ChIP-seq experiments against hypophosphorylated (i.e. PIC/initiation) and total RNA Pol II in NPC derived from our transgenic cell lines. Briefly, these ChIP-seq profiles allowed us to measure (i) PIC formation/initiation (i.e. levels of hypophosphorylated Pol II detected around the *Gata6* TSS) (ii) elongation (i.e. levels of total RNA Pol II detected within the *Gata6* gene body) and (iii) RNA Pol II pause release (i.e. Pausing Index (PI)) for *Gata6* in NPC (see Methods for details). Please note that the antibody against total Pol II also detects an abortive form of RNA Pol II that accumulates around the promoter regions of developmental genes (e.g. *Gata6*) covered by PcG domains (Brookes and Pombo, 2009; Ferrai et al., 2017). In comparison to WT cells, the “core” enhancer alone strongly increased the recruitment of hypophosphorylated Pol II around the *Gata6* TSS (*Enh*: 4.92 ± 0 fold-change over WT) and the insertion site (*Enh*: 6.64 ± 0.23 fold-change over WT) (Fig 6A-B). In contrast, the levels of total RNA Pol II within the *Gata6* gene body were only weakly elevated in *Enh* cells (*Enh*: 2.35 ± 0.12 fold-change over WT) (Fig 6C) and, consequently, the PI was higher in *Enh* cells than in WT cells (Fig 6D). These results suggest that the “core” enhancer alone preferentially regulates PIC formation/initiation to drive *Gata6* expression, thus in agreement with previously proposed models for “typical” enhancers (Larke et al., 2021). Interestingly, the 4xEnh cluster increased hypophosphorylated Pol II levels around the *Gata6* TSS to a similar extent as the “core” enhancer alone (*4xEnh*: 7.13 ± 1.81 fold-change over WT) (Fig 6A-B), but caused a dramatic increase in total RNA Pol II within the *Gata6* gene body (*4xEnh*: 17.13 ± 2.27 fold-change over WT) (Fig 6C). Consequently, the *4xEnh* cells showed an ∼80% reduction in the PI compared to *Enh* cells (Fig 6D). Overall, these results indicate that enhancer clustering synergistically increases *Gata6* expression by preferentially regulating Pol II pause release and productive elongation rather than PIC formation/initiation, which agrees with previous observations for SEs in *Drosophila* (Henriques et al., 2018).

**Fig 6:**
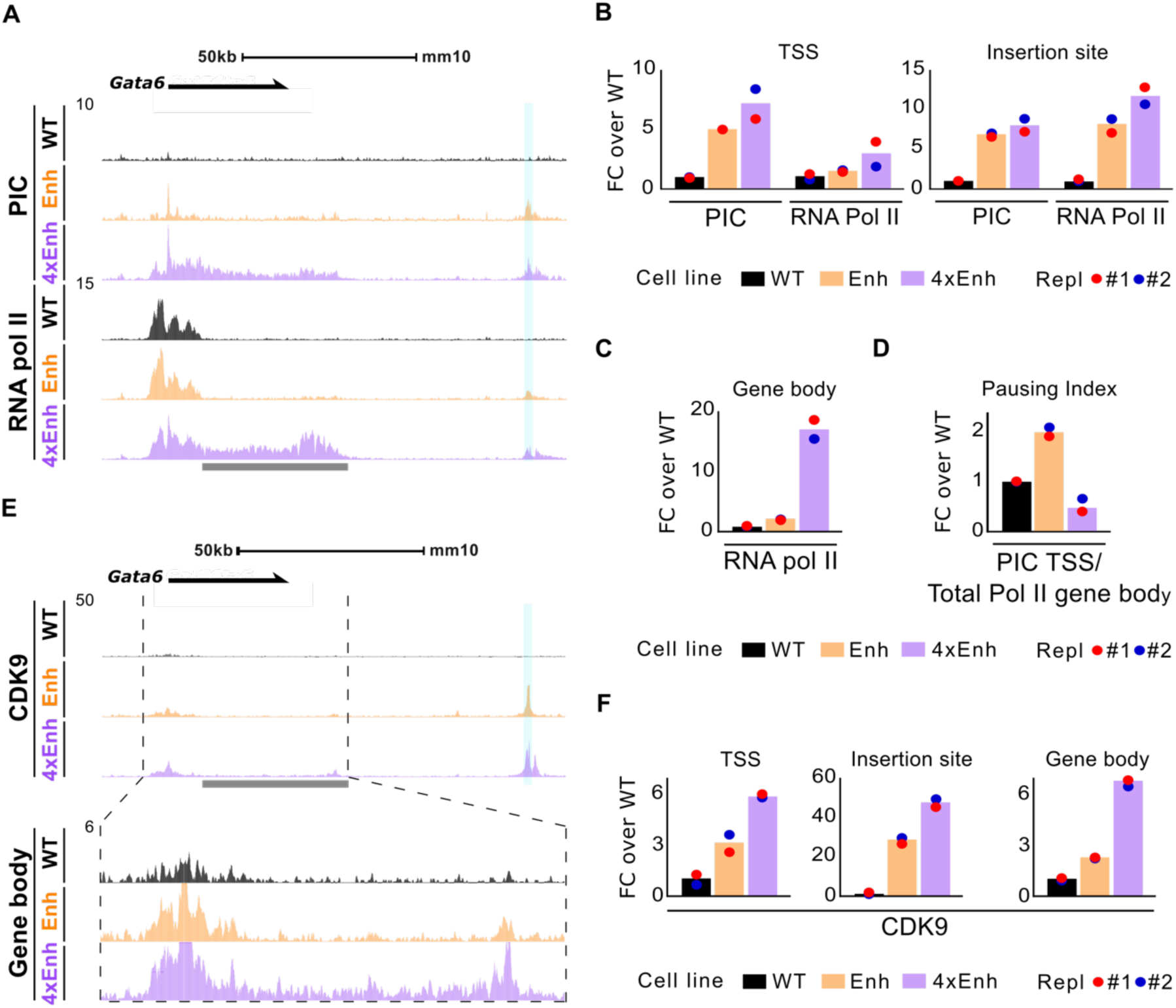
The 4xEnh cluster preferentially controls RNA Pol II pause release and productive transcription rather than PIC formation/initiation. **(A)** ChiP-seq profiles for hypophosphorylated RNA pol II (PIC) and total RNA polymerase II (RNA Pol II) at the Gata6 locus are shown in WT, Enh and 4xEnh NPC. The ChIP-seq tracks represent the average signals from two independent replicates (N=2) for each cell line. The insertion site is highlighted in blue. **(B)** Quantification of the hypophosphorylated RNA pol II (PIC) and total RNA Pol II (RNA Pol II) ChIP-seq signals at the Gata6 transcription start site (TSS) and the synthetic enhancer insertion site in WT, Enh and 4xEnh NPC. For the TSS, the ChIP-seq signals were measured within a +/- 100 bp window centered on the Gata6 TSS. For the insertion site, the ChIP-seq signals were quantified within a region spanning from 1 Kb upstream to 3.5 Kb downstream of the insertion site (Supplementary Data 1). **(C)** Quantification of total RNA Pol II signals across the Gata6 gene body in WT, Enh and 4xEnh NPC. The exact region within which total RNA Pol II signals were quantified is indicated by the gray bar located beneath the ChIP-seq tracks in panel (A) (Supplementary Data 1). ChIP-seq signals within the first two exons were excluded due to the accumulation of abortive RNA Pol II often observed at the 5′ end of Polycomb-bound developmental genes (Ferrai et al., 2017). **(D)** Pausing Indexes (PI) were calculated as the ratio between the PIC signals at the Gata6 TSS (quantified in (B)) and the total RNA pol II signals at the Gata6 gene body (quantified in (C)) (Supplementary Data 1). **(E)** Genome browser tracks showing CDK9 ChIP-seq profiles at the Gata6 locus in WT, Enh and 4xEnh NPC. The ChIP-seq tracks represent the average signals from two independent replicates (N = 2) for each cell line. The region between the vertical dashed lines is highlighted below for better visualization of CDK9 signals within the Gata6 gene body. **(F)** Quantification of CDK9 ChIP-seq signals at the Gata6 TSS, the insertion site and the Gata6 gene body was calculated as described for panels (B) and (C) (Supplementary Data 1). In (B), (C), (D) and (F), the quantifications of the ChIP-seq signals in the synthetic enhancer cell lines are presented as fold-change relative to the ChIP-seq signals in WT cells.

To further explore the dramatic effect that the *4xEnh* cluster had on RNA Pol II pause release and productive elongation, we performed ChIP-seq experiments for CDK9, the catalytic subunit of the pTEBb complex, a key facilitator of Pol II pause release and elongation (Fujinaga et al., 2023). CDK9 is responsible for the phosphorylation of multiple components of the RNA Pol II elongation complex, including serine 2 within the RNA Pol II CTD, and has been shown to travel with the elongating RNA pol II along the body of transcribed genes (Fujinaga et al., 2023). In agreement with the sub-additive increase in the recruitment of several co-activators observed for the 4xEnh cluster, CDK9 levels at the insertion site were almost 2-fold higher in *4xEnh* cells than in *Enh* cells (*Enh*: 26.70 ± 1.29 fold-change over WT; *4xEnh*: 46.34 ± 2.25 fold-change over WT) (Fig 6E-F). More importantly, and in agreement with our results for RNA Pol II, the 4xEnh cluster increased the levels of CDK9 within the *Gata6* gene body compared to the “core” enhancer alone (∼3-fold), while having a more moderate effect at the *Gata6* TSS (2-fold) (Fig 6E-F).

It is worth highlighting that the elevated levels of total RNA Pol II that the *4xEnh* cells show within the *Gata6* gene body are in agreement with the increased contact frequencies observed between the 4xEnh cluster, the *Gata6* promoter and the *Gata6* gene body in those same cells according to the Capture-C experiments described in previous sections (Fig 3B, Fig S2A, Fig S4). Altogether, our smRNA FISH (Fig 2), Capture-C (Fig 3) and RNA Pol II ChIP-seq results (Fig 6) strongly suggest that the 4xEnh cluster increases the proportion of productive enhancer-promoter contacts by facilitating RNA Pol II pause release and the transition into elongation.

## Discussion

Vertebrate genomes utilize a large and heterogeneous repertoire of enhancers to achieve a precise and robust control of developmental gene expression (Boija et al., 2018, 2018; Crispatzu et al., 2021; Kubo et al., 2021; Pachano et al., 2021; Ren et al., 2017; Rinzema et al., 2022; Whyte et al., 2013). Here, we used a synthetic engineering approach to compare the regulatory properties of enhancers with different genetic architectures (i.e. “typical” enhancer, CTCF-associated enhancer and enhancer cluster (resembling a “super-enhancer”)) under identical conditions. This approach minimizes potential confounding factors (e.g. model organism, cell type, type of target promoter, linear distance to target gene, *etc*.) present in previous studies that could restrain accurate comparisons between different enhancer types. Using this approach we showed that (i) the intra-TAD CBS associated with the “core” enhancer mainly works as an insulator facilitating the formation of a chromatin hub/sub-TAD, while having a rather minor effect on enhancer-promoter contacts and target gene expression; (ii) the emergent regulatory properties of enhancer clustering can not be explained by changes in enhancer-promoter contact frequency or the formation of transcriptional condensates. Instead, our data suggest that enhancer clustering synergistically augments target gene expression by preferentially increasing the proportion of transcriptionally productive enhancer–promoter contacts.

Besides the well establish role of CTCF as an insulator at TAD boundaries (Ealo et al., 2024; Nora et al., 2017; Ong and Corces, 2014), CBS are also found within TADs. These intra-TAD CBS are often associated with both enhancers and gene promoters (Kubo et al., 2021; Ren et al., 2017), suggesting an additional role for CTCF as a facilitator of enhancer-gene communication (Rinzema et al., 2022). However, the role of intra-TAD CBS as enhancer tethering elements remains controversial, as the deletion of CBS located nearby enhancers and/or promoters affected target gene expression in some loci (Bower and Kvon, 2025; de Wit et al., 2015; Guo et al., 2015; Ren et al., 2017), but not in others (Chakraborty et al., 2023; Ealo et al., 2024; Pachano et al., 2021). Here we showed that the addition of a CBS to the “core“ enhancer introduced within the *Gata6*-TAD stalls cohesin and generates a smaller sub- domain/chromatin hub (Fig 1E and Fig 3A-C). However, although the interactions amongst genomic regions located inside the newly generated sub-domain were favored, the effects on enhancer-promoter contacts (Fig 3) and, most importantly, on *Gata6* expression were rather mild (Fig 2A). Our findings are in agreement with a recent study in which the effects of CTCF depletion on the 3D chromatin organization and transcriptional activity of 53 different loci were investigated at high resolution during lymphoid-to-myeloid differentiation (Karpinska et al., 2025). We speculate that the previously proposed role for CTCF as a facilitator of enhancer-gene communication might be limited to or especially important for ultra long-range enhancers separated by large linear distances (>400 Kb) from their target genes (Bower and Kvon, 2025). In agreement with this notion, previous studies showed that cohesin-mediated loop extrusion is required for long-range enhancer function (Kane et al., 2022; Rinzema et al., 2022), while short-range enhancers might rely on alternative mechanisms to communicate with their target genes. In this regard, chromatin is subject to passive 3D diffusion motion, which, together with homotypic chromatin interactions, might be sufficient to bring short-range enhancers in proximity of their target genes (Marshall et al., 1997). However, as passive 3D diffusion is inefficient over long linear distances, long-range enhancer-promoter contacts might require cohesin and CTCF assistance (Yang and Hansen, 2024).

Transcription is a rather stochastic and discontinuous process that occurs in bursts (Bahar Halpern et al., 2015; Chubb et al., 2006; Femino et al., 1998; Raj et al., 2006). Previous work suggests that enhancers preferentially control burst frequency/fraction (Bartman et al., 2016; Fukaya et al., 2016; Larson et al., 2013; Larsson et al., 2019; Walters et al., 1995), while burst size is mainly regulated by promoters (Fukaya et al., 2016; Larsson et al., 2019). The effect of enhancers on burst frequency/fraction has been largely attributed to the frequency whereby enhancers contact with their target promoters (Bartman et al., 2016; Tünnermann et al., 2025; Zuin et al., 2022). Interestingly, enhancer-promoter contacts and transcriptional bursts seem to occur with different temporal dynamics, with contacts occurring significantly more often than bursts and, thus, suggesting that only a fraction of all enhancer-promoter contacts might be transcriptionally productive (Cheng et al., 2023; Mach et al., 2022; Tünnermann et al., 2025). Furthermore, live-imaging in cell lines in which a model enhancer was integrated at different linear distances from a reporter gene showed that short-range enhancers not only can drive more frequent but also more uniform bursts than long-range enhancers (Tünnermann et al., 2025). Consequently, it has been suggested that proximal enhancers might confer their target genes with precise and robust expression patterns, while long-range enhancer might lead to more heterogeneous and noisy gene expression. However, this notion is difficult to reconcile with the fact that major developmental genes, which need to be expressed with exquisite precision and specificity, as even subtle changes in their expression can lead to severe congenital defects (Zug, 2022), are often regulated by (ultra) long-range enhancers (Bower and Kvon, 2025). Therefore, long-range enhancers associated with major developmental genes should have unique genetic features allowing these genes to be expressed with high transcriptional precision. Previous studies indicate that certain enhancers might achieve this through nearby tethering elements (e.g. range extender (REX) elements, CpG islands, GAGA-rich elements) that increase the contact frequency with their target promoters over long-distances (Bower et al., 2025; Bower and Kvon, 2025; Cruz-Molina et al., 2017; Li et al., 2023; Li and Levine, 2024; Pachano et al., 2021). Chiefly, our data suggest that enhancer clusters (e.g. super-enhancers) synergistically increase the expression and burst fraction of their target genes, and thereby transcriptional precision, by increasing the proportion of transcriptionally productive enhancer-promoter contacts rather than the frequency of those contacts. More generally, our findings suggest the previously unrecognized possibility that long-range gene expression control might be achieved by modulating the fraction of transcriptionally productive enhancer-promoter contacts. Lastly, considering the complexity of the regulatory landscapes associated with developmental genes, we speculate that the transcriptional precision and robustness that these genes often display might ultimately require the deployment of enhancers with mixed genetic architectures (e.g. super-enhancers linked to tethering elements) (Huang et al., 2018; Ing-Simmons et al., 2015).

Recent models of enhancer function propose that enhancer-promoter spatial proximity might not be required (e.g. TAG model) or could even decrease as enhancers and their target genes become activated (Alexander et al., 2019; Kane et al., 2022; Karr et al., 2022). In this regard, it is worth mentioning that 3C-based methods, such as Capture-C, do not only detect productive but also non-productive enhancer-promoter contacts, as illustrated for example by the pre-formed interactions between primed/poised enhancers and target promoters that precede gene induction in various model organisms (Andrey and Mundlos, 2017; Cruz-Molina et al., 2017; Ghavi-Helm et al., 2014). Therefore, the 4xEnh cluster might not necessarily increase target gene expression by boosting the productivity of enhancer-promoter contacts, as we propose, but through alternative mechanisms that might not even require spatial proximity between enhancers and promoters (e.g. diffusion of activated TFs as proposed by the TAG model; formation of transcriptional condensates). However, the increase in the proportion of productive enhancer-promoter contacts driven by enhancer clustering is further supported by the observed interactions between the 4xEnh cluster, the *Gata6* promoter and the *Gata6* gene body (Fig 3A, B and Fig S2 A, B). These nested interactions are in agreement with previous reports suggesting that, as genes undergo elongation (i.e. productive transcription), enhancers and target promoters track together with the elongating RNA Pol II throughout gene bodies (Lee et al., 2015). Nevertheless, live-imaging approaches enabling the simultaneous visualization of enhancer clusters, target genes and nascent transcripts would be required to distinguish between these alternative models. As mentioned above, one of the mechanisms that could potentially explain how enhancer clusters synergistically increase target gene expression without affecting enhancer-promoter contact frequency is by promoting the formation of transcriptional condensates (Boija et al., 2018; Sabari et al., 2018). These condensates are quite large, with an estimated diameter of ∼500nm, and they might stimulate the expression of a target gene located up to 1um away (Du et al., 2024). As these distances are considerably larger than the ones separating enhancers and promoters engaged into contacts as measured by microscopy or 3C-based methods (<200 nm) (McCord et al., 2020), the nucleation of condensates by enhancer clusters could explain how they synergistically increase target gene expression without altering enhancer-promoter contacts (Yang and Hansen, 2024). Using ChIP-seq, smRNA-FISH and IF assays, we showed that our synthetic 4xEnh cluster displays molecular features typical of SEs, including robust recruitment of Mediator, high levels of H3K27ac and stronger overlap with BRD4 condensates in comparison to the “core” enhancer (Fig 4A-C and Fig 5A-C). However, we found a weak correlation between the levels of BRD4 and *Gata6* nascent transcription in cells with either the “core” enhancer or the *4xEnh* cluster, thus indicating that the sole formation of transcriptional condensates can not explain the emergent regulatory properties and strong regulatory capacity of the enhancer cluster. In this regard, although there are abundant evidences supporting that enhancer and TF clustering promotes the nucleation of transcriptional condensates (Wu et al., 2022), the evidences supporting the direct causal involvement of these condensates in driving gene activation and transcriptional bursts are more scarce (Du et al., 2024; Kawasaki and Fukaya, 2023; Meeussen et al., 2023; Trojanowski et al., 2022; Wu et al., 2022). In fact, recent reports indicate that the strong regulatory activity of IDR-rich TFs are not explained by their propensity to form condensates through LLPS, but rather by their capacity to establish multivalent interactions with multiple co-activators (Meeussen et al., 2023; Trojanowski et al., 2022). Nevertheless, transcriptional condensates display additional features that prevent us from completely ruling out their importance for enhancer cluster function. Namely, as genes sequentially progress through major transcriptional steps (e.g. PIC, initiation, elongation), they might rapidly transition between different types of condensates with mutually exclusively components (Palacio and Taatjes, 2022) (i.e. PIC/initiation condensates rich in Mediator, BRD4 and TFs *vs* elongation and splicing condensates). Consequently, since the 4xEnh cluster facilitates RNA Pol II pause release and elongation, the residence time of *Gata6* and the 4xEnh cluster within PIC/initation condensates might be rather short, rapidly transitioning into elongation/splicing condensates. This could potentially explain the weak correlation between *Gata6* nascent transcription and BRD4 levels.

In addition to facilitating the formation of transcriptional condensates, the co- activators showing strong recruitment to the 4xEnh cluster (i.e. Mediator, p300, BRD4, CDK9) have been previously implicated in the control of several steps along the transcription cycle. Previous work indicates that “typical” enhancers preferentially control PIC formation and/or initiation (Larke et al., 2021). In agreement with this, our RNA Pol II ChIP-seq profiles showed that the “core” enhancer strongly increased PIC formation/initiation in comparison to WT cells (Fig 6A-B). In contrast, the 4xEnh cluster synergistically increased RNA Pol II pause release (i.e. reduced pausing index) and elongation in comparison to the “core” enhancer alone (Fig 6D), while having a moderate effect in PIC formation/initiation. Therefore, our data suggest that enhancer clustering synergistically increases target gene expression by facilitating RNA Pol II pause release rather than PIC formation and/or initiation. Interestingly, this preferential control of pause release by enhancer clusters might be evolutionary conserved, as it was previously reported that SEs in *Drosophila* confer high gene expression levels by enabling rapid pause release (Henriques et al., 2018). Notably, recent studies in vertebrate cells indicate that RNA Pol II displays higher turnover at paused promoters than previously anticipated, suggesting that promoter-proximal RNA Pol II termination is an important and overlooked regulatory step of vertebrate gene transcription (Krebs et al., 2017; Lysakovskaia et al., 2025). As our experimental approach (i.e. ChIP-seq against hypophosphorylated and total RNA Pol II) can not distinguish between polymerases undergoing stable stalling or premature termination at promoter regions, it is possible that enhancer clustering not only increases target gene transcription by facilitating pause release, but also (or even preferentially) by preventing promoter-proximal termination. Regardless, our data strongly supports that RNA Pol II pausing/termination at promoter regions, rather that RNA Pol II recruitment, is the main rate-limiting step that long-range enhancers need to overcome in order to achieve high and precise target gene transcription (Bartman et al., 2019). Intriguingly, for all the investigated co-activators, our ChIP-seq experiments showed that the 4xEnh cluster increased their recruitment in a sub-additive manner, which, nevertheless, resulted in a synergistic transcriptional response. This behavior resembles the multistep ultrasensitivity and bistability associated with signaling cascades, in which a small change in input signal (e.g. overall co-activator levels) can affect multiple steps in a cascade (e.g. transcriptional steps), leading to output amplification (Zhang et al., 2013). Altogether, we propose that a significant proportion of the contacts established between “core” enhancers and their target promoters are transcriptionally non-productive (i.e. contacts that do not result in a transcription burst), because the co- activator levels provided by the enhancer once it is close to its target promoter might be sufficient for PIC formation/initiation but not to overcome RNA Pol II pausing and/or termination. In contrast, as enhancer clusters recruit higher levels of multiple co- activators, upon establishment of an enhancer-promoter contact, this might confer the target promoters with co-activator levels beyond the threshold required for efficient pause release and/or to overcome promoter-proximal termination, thus increasing the proportion of productive enhancer-promoter contacts (i.e. contacts that result in a transcription burst).

## Acknowledgements

We would like to thank all members of the Rada-Iglesias laboratory for their insightful comments and suggestions. Work in the Rada-Iglesias laboratory is supported by the following grants: (i) PID2021- 123030NB-I00, funded by MCIN/AEI/10.13039/501100011033 and by ERDF “A way of making Europe”; (ii) RED2022-134100-T (REDEVNEURAL 3.0), funded by MCIN/AEI/10.13039/501100011033; (iii) ERC Consolidator Grant Poised- Logic (862022), funded by the European Research Council; (iv) ENHPATHY H2020-MSCA-ITN-2019-860002, funded by the European Commission; (v) ChromRare HORIZON-MSCA-2021-DN-01-101073334, funded by the European Commission.

## Materials and Methods

### Cell culture and differentiation

Mouse ESCs (E14Tg2a) were cultured on gelatin-coated plates at 37°C with 5% CO2 in the presence of Knock-out DMEM (Life Technologies, 10829018) supplemented with 15% FBS (Life Technologies, 10082147), 0.1 mM 2-mercaptoethanol (Gibco 21985023), 1x Glutamax (Gibco, 35050061), 1× penicillin–streptomycin (Gibco, 11548876), 1× MEM non-essential amino acids (Gibco, 11140050) and leukemia inhibitory factor (LIF). Differentiation towards Neural Progenitor cells was performed by plating mESC at 13,000 cells/cm2 on geltrex-coated (Gibco, A1413302) plates, which were then grown for six days in N2B27 medium. N2B27 medium was prepared with Advanced Dulbecco’s Modified Eagle Medium F12 (Life Technologies, 21041025) and Neurobasal medium (Life Technologies, 12348017) (1:1), and supplemented with 1× N2 (Life Technologies, 17502048), 1× B27 (Life Technologies, 12587010), 2 mM L-glutamine (Life Technologies, 25030024), 40 mg/ml BSA (Life Technologies, 15260037), 0.1 mM 2-mercaptoethanol and 1× penicillin–streptomycin. During the first three days of differentiation, the N2B27 medium was additionally supplemented with 10 ng/ml bFGF (Life Technologies, PHG0368), while Xav939 (Sigma, 284028-89-3) was added during the last three days at 5 uM final concentration.

### Generation of genetically modified cell lines

The generation of transgenic mouse ESC lines was performed using a gRNAs-Cas9 expressing vector (pX330-hCas9-long-chimeric-grna-g2p; Leo Kurian’s laboratory) and an HDR-template. The HDR templates were generated using a cassette vector in which the synthetic enhancers (Enh-CBS and 4xEnh) were cloned between homology arms flanking the insertion site located 100Kb downstream of *Gata6* (Pachano et al., 2021). *In vitro* digestion of the cassette vectors was performed to isolate the HDR-template consisting of the corresponding synthetic enhancer flanked by the homology arms. The resulting digestion products were resolved on an agarose gel and the product corresponding to the HDR-template was gel-purified (28104, QIAgen). The sequences of the HDR-templates and of the gRNA targeting the insertion site within the *Gata6* locus are shown in Supplementary Data 1.

mESCs were plated at 18,000 cells/cm^2^ on gelatin-coated plates and transfected with Lipofectamine (Thermo Scientific, L3000001) in the presence of the gRNAs-Cas9 expressing vector (250ng) and the corresponding HDR-template (1:1). Cells were cultured in the presence of the transfection reaction in Knock-out DMEM (Life Technologies, 10829018) for 16 hours. Next, cells were grown in Knock out DMEM supplemented with puromycin for 48 hours. Surviving cells were seeded at clonal density in 96-well plates by serial dilution. Clonal mESC lines were genotyped by PCR using the primers listed in Supplementary Data 1. The presence of the modified allele in the derived clones was confirmed by Sanger sequencing.

### RNA extraction and gene expression analysis by RT-qPCR

Total RNA extraction was performed with the NZY total RNA isolation kit (NZYtech) following the manufacturer’s recommendations. ProtoScript II First Strand cDNA Synthesis Kit (New England Biolabs, E6560L) with 1ug of total RNA and random primers was used for cDNA synthesis. RT-qPCRs were performed in technical triplicates for each cDNA samples using the CFX Opus 384 qPCR system (Biorad) with the primers listed in Supplementary Data 1. *Eef1a1* and *Hprt* were used as housekeeping genes.

### smRNA-FISH

smRNA FISH was performed following Stellaris RNA FISH protocol for adherent cells. Briefly, mESC were differentiated into NPC in coverslips coated with geltrex (Gibco, A1413302) at 13,000 cells/cm^2^. At day 6, cells were washed twice with PBS and fixed in 4% paraformaldehyde (VWR, BT140770) in PBS for 10 minutes. Next, cells were washed with PBS twice and incubated overnight in 70% ethanol at 4°C. The next day, cells were incubated at room temperature for 5 minutes in wash buffer A containing 2 x SSC (Invitrogen™ SSC (20X)- AM9770), 10% deionize formamide (Invitrogen Ambion Formamide (Deionized)-AM9342) in RNA-se free water (Invitrogen Ambion Nuclease Free Water- AM9930). The coverslips were placed in a humid chamber on top of a piece of parafilm with a 50ul drop of hybridization buffer containing the probe and incubated overnight at 37°C. Hybridization buffer: 10% dextran sulfate sodium salt (Sigma Aldrich-D8906-5G), 10% mL deionized formamide (Invitrogen Ambion Formamide (Deionized)-AM9342), 2 x SSC (Invitrogen™ SSC (20X)- AM9770) in 8 mL RNA-free water (Invitrogen Ambion Nuclease Free Water- AM9930) and probe (12.5mM RNA). Afterwards, cells were incubated for 30 min in wash buffer B (2xSCC in RNA-free water) at 37°C. Next, an additional incubation with wash buffer B containing DAPI (Sigma-aldrich- D9542-1MG) was performed. Finally, cells were washed for 5 min in wash buffer B at room temperature and the coverslips were mounted onto glass slides with Vectashield (VWR, 101098-042) and sealed with nail polish for subsequent image acquisition. Intronic and exonic probes were designed using the Stellaris Probe Designer (https://www.biosearchtech.com/support/tools/design-software/stellaris-probe-designer) following manufacturer’s recommendations and are listed in Supplementary Data 1.

### Combination of immunofluorescence and smRNA-FISH assays

IF assays were combined with smRNA-FISH as described in (Boija et al., 2018; Sabari et al., 2018) with minor modifications. Briefly, mESC were plated in coverslips coated with geltrex (Gibco, A1413302) at 37°C and differentiated in the presence of N2B27. At day 6, cells were washed twice with PBS and fixed for 10 minutes using 4% paraformaldehyde (VWR, BT140770) in PBS. After fixation, cells were washed twice with PBS and incubated overnight in 70% ethanol at 4C. The next day, cells were washed twice with PBS and permeabilized with 0.5% Triton X-100 (Sigma Aldrich, X100) in PBS for 10 min, followed by two additional washes in PBS. Next, cells were incubated for 30 min in 4% IgG-free Bovine Serum Albumin (BSA; VWR, 102643-516) and then overnight at 4°C with the primary antibody against BRD4 (ab128874) at a 1:500 concentration in PBS. The next day, cells were washed three times with PBS and incubated with secondary antibody (Invitrogen, A32731) at a 1:5000 concentration in PBS for 1 hour. After washing three times with PBS, cells were fixed for 10 minutes with 4% paraformaldehyde (PFA; VWR, BT140770) in PBS. After washing twice with PBS, smRNA-FISH was performed as described above.

### Image acquisition and analyses

Images were acquired on a NIKON Eclipse Ti2 fluorescence microscope with a 100x oil objective using NIS-elements acquisition software and the ORCA Flash4.0 (Hamamatsu) camera. Images were post-processed using Fiji Is Just ImageJ (FIJI).

Nascent transcription foci were identified in 3D by generating maximum intensity Z-stack projections. To minimize background and out-of-focus signal, the first slice of each image stack was discarded, and projections were generated using the subsequent five slices, acquired at 0.5 μm intervals. Next, channels were split, and a Gaussian blur filter (σ = 1) was applied, followed by threshold adjustment and particle analysis (size: 50 µm² to ∞) to exclude background and saturated regions. After performing maxima projection of the different Z-stack for the different channels a 9×9 convolution kernel was applied to enhance foci contrast across channels. Exonic and intronic background noise was estimated by averaging the standard deviation of three randomly selected 20×20 µm² regions in each channel. Maxima corresponding to exonic and intronic signal were identified using a prominence threshold set to ten times the standard deviation. To refine the overlap between intronic and exonic maxima, a binary dilation was applied to single-point maxima identified in each channel. Overlapping maxima were used to define nascent transcription foci. Identified foci were manually curated for validation. Next, the intensity of the defined intronic signal was measured from raw Z-stack projections to determine burst size. To correct for local background, the signal surrounding each identified intronic maximum was measured and subtracted from the corresponding maximum. Analyses were performed in two independent replicates for each genotype. For burst fraction quantification, the number of transcriptionally active alleles were assessed in at least 1,000 cells across two independent replicates.

### Cumulative plots for *Gata6* smRNA FISH and BRD4 IF signals

For smRNA-FISH combined with IF against BRD4, the IF signals were centered on manually selected nascent FISH foci and averaged to generate a composite profile of BRD4 enrichment at *Gata6* nascent transcription sites (i.e. foci with colocalizing intronic and exonic signals). One hundred *Gata6* nascent FISH foci were selected from two independent replicates per genotype. For each nascent transcription site selected, the Z-slice with the highest intronic signal was manually identified and the two preceding and two following slices, acquired at 0.5 μm intervals, were included to quantify BRD4 signals around the transcription site. The selected Z-slices from the 100 foci were concatenated to create a unified dataset. To improve the definition of both FISH and IF puncta, a Gaussian blur filter (σ = 1) was applied, followed by radial background subtraction (σ = 10). Next, the aggregated signals for both nascent FISH and BRD4 were obtained by generating maximum intensity Z-stack projections across the selected slices, with BRD4 signals displayed as a contour map. Intronic and BRD4 signals were quantified within a 3 µm³ volume centered on the intronic signal maximum for each defined transcription site. This region encompassed a 3 µm diameter area in the XY plane and spanned three consecutive Z-slices (±1 slice from the focal plane). As a control, the same process was carried out for IF signals centered at 200 randomly selected nuclear positions for each genotype. For correlation analyses, the IF signals on the 200 randomly selected regions (i.e. background regions) were subtracted from the IF signals quantified at the *Gata6* nascent transcription sites.

### Capture-C

Capture-C was performed in two biological replicates per genotype as previously described (Oudelaar et al., 2018). Briefly, 5 × 10^6^ cells were crosslinked with 2% paraformaldehyde for 10 min. Subsequently, cells were quenched with 0.125 M glycine for 10 min. After washing three times with PBS, cells were resuspended in lysis buffer (10 mM Tris pH 8, 10 mM NaCl, 0.2% NP-40 and 1× protease inhibitors) for 30 min on ice. Afterwards, nuclei were pelleted by centrifugation and resuspended in NEB Buffer DpnII (B0543S, NEB) for subsequent chromatin digestion with DpnII (R0543S, NEB) followed by proximity ligation (Life Tech, cat. no. EL0013). Next, de-crosslinking was performed with Proteinase K (3U; Thermo Fisher, cat. no. EO0491) and subsequent incubation with RNase A (7.5 mU; Roche: 1119915). Afterwards, DNA was purified using the Qiagen kit (28506). Purified DNA was sonicated using Covaris S220 Focused-ultrasonicator (time, 180 s; duty factor, 10%; peak incident power, 175 W; cycles per burst, 200) to an average size of 200 bp. Sheared DNA was purified using AMPure XP SPRI beads (Beckman Coulter, cat. no. A63881). 3C libraries were generated using the NEBNext Ultra II kit (New England Biolabs, cat. no. E7645S/L). Index primers set 1 and 2 from the NEBNext Multiplex Oligos for Illumina kit (New England Biolabs, E7335S/L E7500S/L) were incorporated using Herculase II Fusion Polymerase Kit (Agilent, cat. no. 600677). Enrichment of ligated fragments containing the viewpoints of interest was performed using 120bp long probes in two consecutive rounds, as described for MCC(Hua et al., 2021). Isolation of hybridized DNA was performed using M-270 Streptavidin Dynabeads (Invitrogen, 65305) and amplified by PCR (11 cycles). The second enrichment reaction was performed using the pulled down material from the first enrichment reaction as input. The quality of the final libraries was assessed with a fragment analyzer and sequenced using the the Novaseq6000 sequencing system (150bp paired-end reads;15 Mreads/lib). The sequences of the different 120 bp probes used for the enrichment reactions (i.e. viewpoints) as shown in Supplementary Data 1.

Capture-C data obtained from the different cell lines was analyzed using the CapCruncher pipeline (version 0.2.3)(Hua et al., 2021) using *mm10* as a reference genome. For the cell lines containing synthetic enhancers, custom genomes were generated using Reform, a Python-based command-line tool (https://gencore.bio.nyu.edu/reform). Custom genomes lack the endogenous genomic sequences used to engineer the synthetic enhancers (i.e. the *Sox1*+35kb “core” enhancer or the CBS at the *Shh* locus) at the endogenous loci and included the corresponding synthetic enhancer at the insertion site.

### Chromatin immunoprecipitation (ChIP)

ChIPs for histone marks were performed using approximately 1 × 10⁷ cells, while ∼4 × 10⁷ cells were used for all other proteins. The antibodies used are listed in Supplementary Data 1. Double crosslinking was performed for MED26 and BRD4 ChIPs. Briefly, cells were washed twice with PBS and fixed for 30 min at room temperature in the presence of ChIP crosslink-Gold (Diagenode, C01019027) following the manufacturer’s recommendations. The subsequent steps were the same as for the remaining ChIPs in which a single crosslinking step was performed. Single crosslinking was performed by initially washing twice with PBS before incubation with 1% formaldehyde (VWR, BT140770) in KO DMEM for 10 min at room temperature (RT). Next, the formaldehyde was quenched with 0.125 M glycine for 10 min. Chromatin isolation was performed by consecutive incubation in three different lysis buffers, each for 10 min (Buffer 1: 50 mM HEPES, 140 mM NaCl, 1 mM EDTA, 10% glycerol, 0.5% NP-40, 0.25% TX-100; Buffer 2: 10 mM Tris, 200 mM NaCl, 1 mM EDTA, 0.5 mM EGTA; Buffer 3: 10 mM Tris, 100 mM NaCl, 1 mM EDTA, 0.5 mM EGTA, 0.1% Na-deoxycholate, 0.5% N-lauroylsarcosine). Chromatin was then sonicated using an EpiShear probe sonicator (Active Motif) for 4 min in Buffer 3 (20 s on, 30 s off, 30% amplitude). Sonicated chromatin was incubated overnight at 4 °C with 5 μg of antibody for the different histone marks or with 10 μg of antibody for the remaining proteins. Then, 50 μl of Dynabeads (Thermo Fisher, 10-002-D) were added and immunoprecipitations were performed for 4 hours at 4 °C. The beads were pre-washed with cold blocking solution (0.5% BSA in 1× PBS). Once the immunoprecipitation was finished, the beads were washed five times with RIPA buffer (50 mM HEPES, 500 mM LiCl, 1 mM EDTA, 1% NP-40, and 0.7% sodium deoxycholate). Chromatin was eluted by incubation of the beads at 65 °C overnight, followed by treatment with RNase A (0.2 mg/ml) for 1 hour at 37 °C, and then with proteinase K (0.2 mg/ml) at 55 °C for 2h. Finally, DNA was purified using the Zymoclean Gel DNA Recovery Kit (Zymo Research, D4008).

### ChIP-seq analyses: Sequencing, alignment, quantification and normalization

Library preparation for paired-end sequencing was performed using TruSeq DNA Sample prep Kit (Illumina) following the manufacturer’s recommendations and sequenced using the Novaseq6000 sequencing system (Illumina; 150bp paired-end reads, 40Mreads/library). Quality control and trimming of low quality reads and/or adapters was performed using *fastqc* (https://www.bioinformatics.babraham.ac.uk/projects/fastqc/), *MultiQC* (Ewels et al., 2016) and *trimmomatic* (Bolger et al., 2014). Reads were subsequently mapped to the mm10 reference genome using Bowtie2 (Langmead and Salzberg, 2012). For transgenic cell lines, custom genomes including the synthetic enhancers at the corresponding loci were generated using *Reform*, a Python-based command-line tool (https://gencore.bio.nyu.edu/reform). Since the synthetic enhancers were designed based on endogenous mouse sequences (specifically the *Sox1+35kb* core enhancer or the CBS at the *Shh* locus), Bowtie2 alignment was restricted to concordant alignments, allowing up to two valid alignments per read. Only reads with mapping quality above 10 were considered for alignments, and duplicated reads were removed with SAMtools (Li et al., 2009). Next, batch effect correction was performed by applying quantile normalization separately for each antibody across the different genotypes and replicates(Mariner-Faulí et al., 2025). Briefly, RPKM values were calculated using 100 bp bins and, subsequently, quantile normalization was applied across all samples. The resulting bedGraph files were used for quantification of ChIP-seq signals in the regions indicated in Supplementary Data 1 using computeMatrix from deepTools (Ramírez et al., 2016). Next, the BedGraph files from the two replicates generated for each sample were averaged and converted to BigWig format using the *bedGraphToBigWig* tool from UCSC (Kent et al., 2010) in order to visualize the different ChIP-seq profiles on the UCSC genome browser (Perez et al., 2025).

For calculating the Pausing Index (PI) for *Gata6*, the amount of ChIP-seq signal for hypophosphorylated RNA pol II around the *Gata6* TSS relative to the amount of ChIP-seq signal for total RNA Pol II within the *Gata6* gene body was calculated. The genomic regions considered for measuring ChIP-seq signals within the *Gata6* TSS and the *Gata6* gene body are indicated in Supplementary Data 1.

### Data availability

All the genomic datasets generated in this project have been deposited into GEO. They will be publicly available upon peer-reviewed publication of this work.

## Supplementary figures

**Fig S1:**
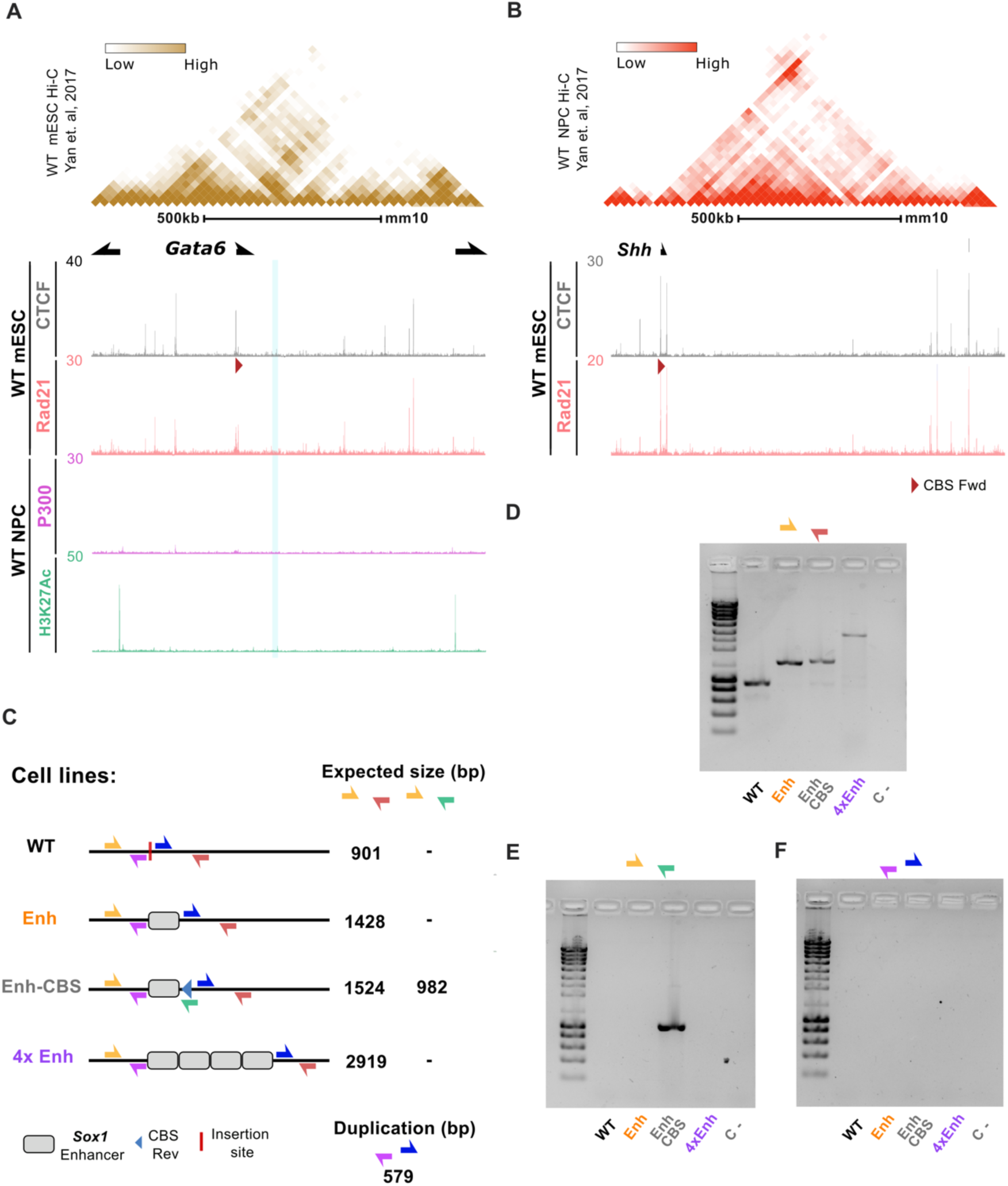
Generation and characterization of transgenic mESC lines with synthetic enhancers inserted within the Gata6 locus. **(A)** mESC Hi-C data at the Gata6 locus (Yan et al., 2018) is shown on top to illustrate the location of the Gata6-TAD. Below, ChIP-seq tracks for CTCF and Rad21 in WT mESC together with ChIP-seq tracks for p300 and H3K27ac in WT NPC are shown. The blue highlighted region indicates the insertion site of the synthetic enhancers. **(B)** Shh TAD in mESC according to previously generated Hi-C data (Yan et al., 2018). Below, CTCF and Rad21 ChIP-seq tracks from WT mESC are shown at the Shh locus, with the CTCF peak containing the CBS used for the generation of the Enh-CBS synthetic enhancer (Ehn-CBS) indicated with a red arrow. **(C)** Genotyping strategy used for the different synthetic enhancers inserted at the Gata6 locus. The expected size for the PCR products is indicated on the right. **(D)** Genotyping results obtained with the PCR primers described in (C). **(E)** Same as in D, but with a different set of primers for the specific amplification of the Enh- CBS insert. **(F)** Same as in D but with primers designed to identify possible duplications.

**Fig S2:**
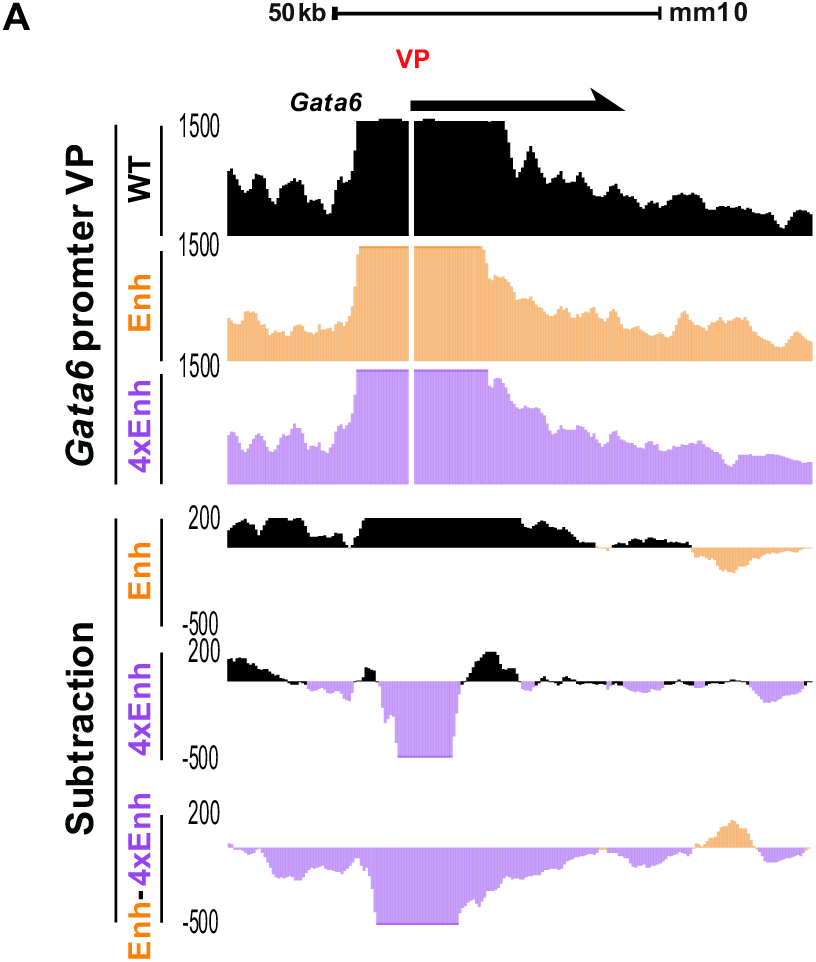
The Gata6 promoter shows increased contact frequency with the Gata6 gene body in 4xEnh cells. Zoomed-in view of the Capture-C signals from Fig 3A are shown around the Gata6 gene body to better visualize the contacts between the Gata6 promoter and the Gata6 gene body. The Capture-C profiles were generated using the Gata6 promoter as a viewpoint (VP) and are shown for NPC derived from the indicated cell lines. For each cell line, the Capture-C profiles were generated by averaging the signals from two independent replicates (N = 2). Underneath, subtraction tracks are presented in order to highlight the differences in Capture-C signals between cell lines.

**Fig S3:**
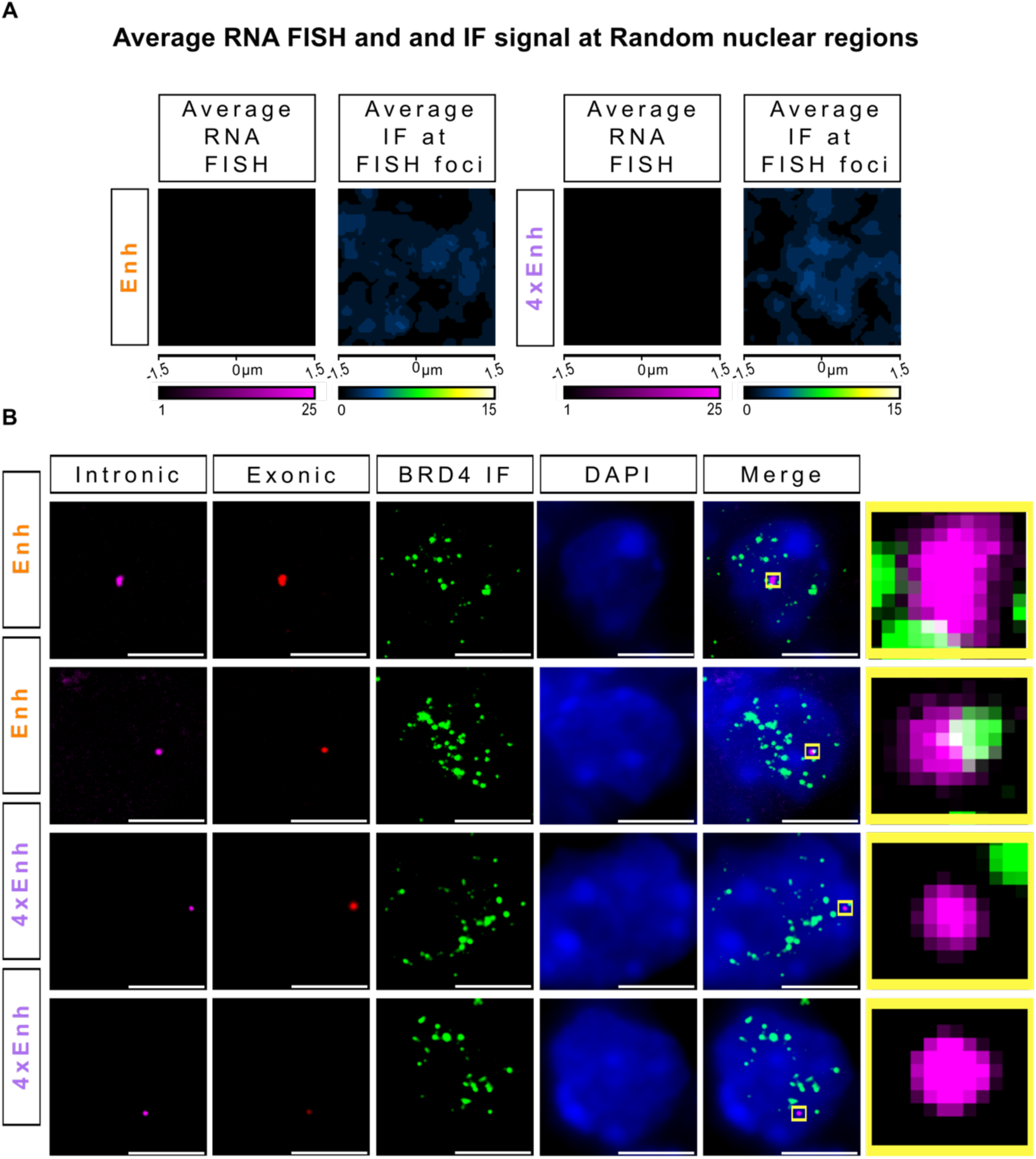
Combination of BRD4 IF and Gata6 smRNA FISH assays in Enh and 4xEnh NPC. **(A)** Average intronic Gata6 smRNA-FISH (left) and BRD4 IF signals (right) centered around 200 random nuclear regions in Enh and 4xEnh NPC using data from two independent replicates per genotype. **(B)** Co-localization of BRD4 IF and Gata6 smRNA-FISH signals within selected cells of the indicated genotypes (i.e. Enh or 4xEnh). Individual panels display the BRD4 signal (green) and single-molecule RNA FISH signals targeting either exonic (red) or intronic (magenta) Gata6 regions in NPC derived from the indicated ESC lines. A composite image (Merge) including DAPI staining is shown to highlight the co-localization of the different signals. A zoomed-in view of the area highlighted by the yellow box is provided for clearer visualization of the overlap between IF and intronic signals. Scale bar: 5 µm.

**Fig S4:**
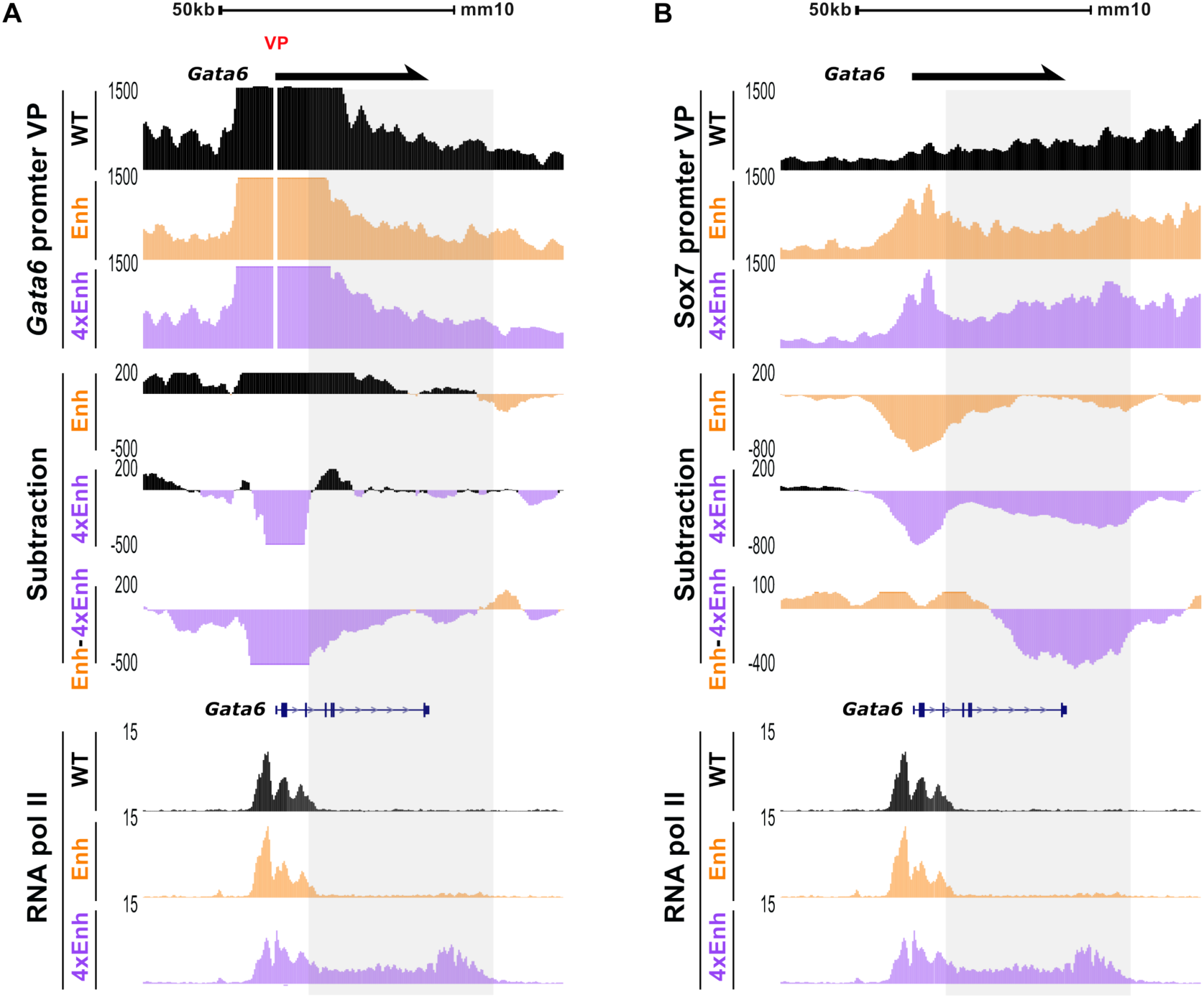
The increased contact frequency between the 4xEnh cluster and the Gata6 promoter with the Gata6 gene body in 4xEnh cells resembles the distribution of elongating RNA pol II. **(A-B)** Zoomed-in view of the Capture-C signals and subtraction tracks from Fig 3A-B around the Gata6 gene body are shown on top for the indicated genotypes using either the Gata6 promoter **(A)** or the insertion site **(B)** as viewpoints. Underneath, total RNA pol II ChIP-seq signals from Fig 6A are shown around the Gata6 gene body. For each cell line, the Capture-C profiles were generated by averaging the signals from two independent replicates (N = 2).

## References

1. Aboreden, N.G., Lam, J.C., Goel, V.Y., Wang, S., Wang, X., Midla, S.C., Quijano, A., Keller, C.A., Giardine, B.M., Hardison, R.C., Zhang, H., Hansen, A.S., Blobel, G.A., 2025. LDB1 establishes multi-enhancer networks to regulate gene expression. Mol. Cell 85, 376–393.e9. 10.1016/j.molcel.2024.11.037

2. Alexander, J.M., Guan, J., Li, B., Maliskova, L., Song, M., Shen, Y., Huang, B., Lomvardas, S., Weiner, O.D., 2019. Live-cell imaging reveals enhancer-dependent Sox2 transcription in the absence of enhancer proximity. eLife 8, e41769. 10.7554/eLife.41769

3. Altendorfer, E., Mochalova, Y., Mayer, A., 2022. BRD4: a general regulator of transcription elongation. Transcription 13, 70–81. 10.1080/21541264.2022.2108302

4. Andrey, G., Mundlos, S., 2017. The three-dimensional genome: regulating gene expression during pluripotency and development. Development 144, 3646–3658. 10.1242/dev.148304

5. Bahar Halpern, K., Tanami, S., Landen, S., Chapal, M., Szlak, L., Hutzler, A., Nizhberg, A., Itzkovitz, S., 2015. Bursty Gene Expression in the Intact Mammalian Liver. Mol. Cell 58, 147–156. 10.1016/j.molcel.2015.01.027

6. Bartman, C.R., Hamagami, N., Keller, C.A., Giardine, B., Hardison, R.C., Blobel, G.A., Raj, A., 2019. Transcriptional Burst Initiation and Polymerase Pause Release Are Key Control Points of Transcriptional Regulation. Mol. Cell 73, 519–532.e4. 10.1016/j.molcel.2018.11.004

7. Bartman, C.R., Hsu, S.C., Hsiung, C.C.-S., Raj, A., Blobel, G.A., 2016. Enhancer Regulation of Transcriptional Bursting Parameters Revealed by Forced Chromatin Looping. Mol. Cell 62, 237–247. 10.1016/j.molcel.2016.03.007

8. Boija, A., Klein, I.A., Sabari, B.R., Dall’Agnese, A., Coffey, E.L., Zamudio, A.V., Li, C.H., Shrinivas, K., Manteiga, J.C., Hannett, N.M., Abraham, B.J., Afeyan, L.K., Guo, Y.E., Rimel, J.K., Fant, C.B., Schuijers, J., Lee, T.I., Taatjes, D.J., Young, R.A., 2018. Transcription Factors Activate Genes through the Phase-Separation Capacity of Their Activation Domains. Cell 175, 1842–1855.e16. 10.1016/j.cell.2018.10.042

9. Bolger, A.M., Lohse, M., Usadel, B., 2014. Trimmomatic: a flexible trimmer for Illumina sequence data. Bioinformatics 30, 2114–2120. 10.1093/bioinformatics/btu170

10. Bower, G., Hollingsworth, E.W., Jacinto, S.H., Alcantara, J.A., Clock, B., Cao, K., Liu, M., Dziulko, A., Alcaina-Caro, A., Xu, Q., Skowronska-Krawczyk, D., Lopez-Rios, J., Dickel, D.E., Bardet, A.F., Pennacchio, L.A., Visel, A., Kvon, E.Z., 2025. Range extender mediates long-distance enhancer activity. Nature 643, 830–838. 10.1038/s41586-025-09221-6

11. Bower, G., Kvon, E.Z., 2025. Genetic factors mediating long-range enhancer–promoter communication in mammalian development. Curr. Opin. Genet. Dev. 90, 102282. 10.1016/j.gde.2024.102282

12. Brookes, E., Pombo, A., 2009. Modifications of RNA polymerase II are pivotal in regulating gene expression states. EMBO Rep. 10, 1213–1219. 10.1038/embor.2009.221

13. Chakraborty, S., Kopitchinski, N., Zuo, Z., Eraso, A., Awasthi, P., Chari, R., Mitra, A., Tobias, I.C., Moorthy, S.D., Dale, R.K., Mitchell, J.A., Petros, T.J., Rocha, P.P., 2023. Enhancer– promoter interactions can bypass CTCF-mediated boundaries and contribute to phenotypic robustness. Nat. Genet. 55, 280–290. 10.1038/s41588-022-01295-6

14. Chen, H., Levo, M., Barinov, L., Fujioka, M., Jaynes, J.B., Gregor, T., 2018. Dynamic interplay between enhancer–promoter topology and gene activity. Nat. Genet. 50, 1296–1303. 10.1038/s41588-018-0175-z

15. Chen, L., Zhang, Z., Han, Q., Maity, B.K., Rodrigues, L., Zboril, E., Adhikari, R., Ko, S.-H., Li, X., Yoshida, S.R., Xue, P., Smith, E., Xu, K., Wang, Q., Huang, T.H.-M., Chong, S., Liu, Z., 2023. Hormone-induced enhancer assembly requires an optimal level of hormone receptor multivalent interactions. Mol. Cell 83, 3438–3456.e12. 10.1016/j.molcel.2023.08.027

16. Chen, Z., Snetkova, V., Bower, G., Jacinto, S., Clock, B., Dizehchi, A., Barozzi, I., Mannion, B.J., Alcaina-Caro, A., Lopez-Rios, J., Dickel, D.E., Visel, A., Pennacchio, L.A., Kvon, E.Z., 2024. Increased enhancer-promoter interactions during developmental enhancer activation in mammals. Nat. Genet. 56, 675–685. 10.1038/s41588-024-01681-2

17. Cheng, L., De, C., Li, J., Pertsinidis, A., 2023. Mechanisms of transcription control by distal enhancers from high-resolution single-gene imaging. BioRxiv Prepr. Serv. Biol. 2023.03.19.533190. 10.1101/2023.03.19.533190

18. Chubb, J.R., Trcek, T., Shenoy, S.M., Singer, R.H., 2006. Transcriptional Pulsing of a Developmental Gene. Curr. Biol. 16, 1018–1025. 10.1016/j.cub.2006.03.092

19. Cramer, P., 2019. Organization and regulation of gene transcription. Nature 573, 45–54. 10.1038/s41586-019-1517-4

20. Crispatzu, G., Rehimi, R., Pachano, T., Bleckwehl, T., Cruz-Molina, S., Xiao, C., Mahabir, E., Bazzi, H., Rada-Iglesias, A., 2021. The chromatin, topological and regulatory properties of pluripotency-associated poised enhancers are conserved in vivo. Nat. Commun. 12, 4344. 10.1038/s41467-021-24641-4

21. Cruz-Molina, S., Respuela, P., Tebartz, C., Kolovos, P., Nikolic, M., Fueyo, R., Van Ijcken, W.F.J., Grosveld, F., Frommolt, P., Bazzi, H., Rada-Iglesias, A., 2017. PRC2 Facilitates the Regulatory Topology Required for Poised Enhancer Function during Pluripotent Stem Cell Differentiation. Cell Stem Cell 20, 689–705.e9. 10.1016/j.stem.2017.02.004

22. Dao, L.T.M., Galindo-Albarrán, A.O., Castro-Mondragon, J.A., Andrieu-Soler, C., Medina-Rivera, A., Souaid, C., Charbonnier, G., Griffon, A., Vanhille, L., Stephen, T., Alomairi, J., Martin, D., Torres, M., Fernandez, N., Soler, E., Van Helden, J., Puthier, D., Spicuglia, S., 2017. Genome-wide characterization of mammalian promoters with distal enhancer functions. Nat. Genet. 49, 1073–1081. 10.1038/ng.3884

23. de Wit, E., Vos, E.S.M., Holwerda, S.J.B., Valdes-Quezada, C., Verstegen, M.J.A.M., Teunissen, H., Splinter, E., Wijchers, P.J., Krijger, P.H.L., de Laat, W., 2015. CTCF Binding Polarity Determines Chromatin Looping. Mol. Cell 60, 676–684. 10.1016/j.molcel.2015.09.023

24. Dixon, J.R., Selvaraj, S., Yue, F., Kim, A., Li, Y., Shen, Y., Hu, M., Liu, J.S., Ren, B., 2012. Topological domains in mammalian genomes identified by analysis of chromatin interactions. Nature 485, 376–380. 10.1038/nature11082

25. Dowen, J.M., Fan, Z.P., Hnisz, D., Ren, G., Abraham, B.J., Zhang, L.N., Weintraub, A.S., Schujiers, J., Lee, T.I., Zhao, K., Young, R.A., 2014. Control of cell identity genes occurs in insulated neighborhoods in mammalian chromosomes. Cell 159, 374–387. 10.1016/j.cell.2014.09.030

26. Downes, D.J., Smith, A.L., Karpinska, M.A., Velychko, T., Rue-Albrecht, K., Sims, D., Milne, T.A., Davies, J.O.J., Oudelaar, A.M., Hughes, J.R., 2022. Capture-C: a modular and flexible approach for high-resolution chromosome conformation capture. Nat. Protoc. 17, 445–475. 10.1038/s41596-021-00651-w

27. Du, M., Stitzinger, S.H., Spille, J.-H., Cho, W.-K., Lee, C., Hijaz, M., Quintana, A., Cissé, I.I., 2024. Direct observation of a condensate effect on super-enhancer controlled gene bursting. Cell 187, 331–344.e17. 10.1016/j.cell.2023.12.005

28. Ealo, T., Sanchez-Gaya, V., Respuela, P., Muñoz-San Martín, M., Martin-Batista, E., Haro, E., Rada-Iglesias, A., 2024. Cooperative insulation of regulatory domains by CTCF-dependent physical insulation and promoter competition. Nat. Commun. 15, 7258. 10.1038/s41467-024-51602-4

29. Ewels, P., Magnusson, M., Lundin, S., Käller, M., 2016. MultiQC: summarize analysis results for multiple tools and samples in a single report. Bioinformatics 32, 3047–3048. 10.1093/bioinformatics/btw354

30. Femino, A.M., Fay, F.S., Fogarty, K., Singer, R.H., 1998. Visualization of Single RNA Transcripts in Situ. Science 280, 585–590. 10.1126/science.280.5363.585

31. Ferrai, C., Torlai Triglia, E., Risner-Janiczek, J.R., Rito, T., Rackham, O.J., De Santiago, I., Kukalev, A., Nicodemi, M., Akalin, A., Li, M., Ungless, M.A., Pombo, A., 2017. RNA polymerase II primes Polycomb-repressed developmental genes throughout terminal neuronal differentiation. Mol. Syst. Biol. 13, 946. 10.15252/msb.20177754

32. Fuda, N.J., Ardehali, M.B., Lis, J.T., 2009. Defining mechanisms that regulate RNA polymerase II transcription in vivo. Nature 461, 186–192. 10.1038/nature08449

33. Fudenberg, G., Imakaev, M., Lu, C., Goloborodko, A., Abdennur, N., Mirny, L.A., 2016. Formation of Chromosomal Domains by Loop Extrusion. Cell Rep. 15, 2038–2049. 10.1016/j.celrep.2016.04.085

34. Fujinaga, K., Huang, F., Peterlin, B.M., 2023. P-TEFb: The master regulator of transcription elongation. Mol. Cell 83, 393–403. 10.1016/j.molcel.2022.12.006

35. Fukaya, T., Lim, B., Levine, M., 2016. Enhancer Control of Transcriptional Bursting. Cell 166, 358–368. 10.1016/j.cell.2016.05.025

36. Ghavi-Helm, Y., Klein, F.A., Pakozdi, T., Ciglar, L., Noordermeer, D., Huber, W., Furlong, E.E.M., 2014. Enhancer loops appear stable during development and are associated with paused polymerase. Nature 512, 96–100. 10.1038/nature13417

37. Grosveld, F., Van Assendelft, G.B., Greaves, D.R., Kollias, G., 1987. Position-independent, high-level expression of the human β-globin gene in transgenic mice. Cell 51, 975–985. 10.1016/0092-8674(87)90584-8

38. Guo, Y., Xu, Q., Canzio, D., Shou, J., Li, J., Gorkin, D.U., Jung, I., Wu, H., Zhai, Y., Tang, Y., Lu, Y., Wu, Y., Jia, Z., Li, W., Zhang, M.Q., Ren, B., Krainer, A.R., Maniatis, T., Wu, Q., 2015. CRISPR Inversion of CTCF Sites Alters Genome Topology and Enhancer/Promoter Function. Cell 162, 900–910. 10.1016/j.cell.2015.07.038

39. Heintzman, N.D., Hon, G.C., Hawkins, R.D., Kheradpour, P., Stark, A., Harp, L.F., Ye, Z., Lee, L.K., Stuart, R.K., Ching, C.W., Ching, K.A., Antosiewicz-Bourget, J.E., Liu, H., Zhang, X., Green, R.D., Lobanenkov, V.V., Stewart, R., Thomson, J.A., Crawford, G.E., Kellis, M., Ren, B., 2009. Histone modifications at human enhancers reflect global cell-type-specific gene expression. Nature 459, 108–112. 10.1038/nature07829

40. Henriques, T., Scruggs, B.S., Inouye, M.O., Muse, G.W., Williams, L.H., Burkholder, A.B., Lavender, C.A., Fargo, D.C., Adelman, K., 2018. Widespread transcriptional pausing and elongation control at enhancers. Genes Dev. 32, 26–41. 10.1101/gad.309351.117

41. Hnisz, D., Abraham, B.J., Lee, T.I., Lau, A., Saint-André, V., Sigova, A.A., Hoke, H.A., Young, R.A., 2013. Super-Enhancers in the Control of Cell Identity and Disease. Cell 155, 934–947. 10.1016/j.cell.2013.09.053

42. Hua, P., Badat, M., Hanssen, L.L.P., Hentges, L.D., Crump, N., Downes, D.J., Jeziorska, D.M., Oudelaar, A.M., Schwessinger, R., Taylor, S., Milne, T.A., Hughes, J.R., Higgs, D.R., Davies, J.O.J., 2021. Defining genome architecture at base-pair resolution. Nature 595, 125–129. 10.1038/s41586-021-03639-4

43. Huang, J., Li, K., Cai, W., Liu, X., Zhang, Y., Orkin, S.H., Xu, J., Yuan, G.-C., 2018. Dissecting super-enhancer hierarchy based on chromatin interactions. Nat. Commun. 9, 943. 10.1038/s41467-018-03279-9

44. Ing-Simmons, E., Seitan, V.C., Faure, A.J., Flicek, P., Carroll, T., Dekker, J., Fisher, A.G., Lenhard, B., Merkenschlager, M., 2015. Spatial enhancer clustering and regulation of enhancer-proximal genes by cohesin. Genome Res. 25, 504–513. 10.1101/gr.184986.114

45. Jiang, Y., Qian, F., Bai, X., Liu, Y., Wang, Q., Ai, B., Han, X., Shi, S., Zhang, J., Li, X., Tang, Z., Pan, Q., Wang, Y., Wang, F., Li, C., 2019. SEdb: a comprehensive human super-enhancer database. Nucleic Acids Res. 47, D235–D243. 10.1093/nar/gky1025

46. Kane, L., Williamson, I., Flyamer, I.M., Kumar, Y., Hill, R.E., Lettice, L.A., Bickmore, W.A., 2022. Cohesin is required for long-range enhancer action at the Shh locus. Nat. Struct. Mol. Biol. 29, 891–897. 10.1038/s41594-022-00821-8

47. Karpinska, M.A., Zhu, Y., Fakhraei Ghazvini, Z., Ramasamy, S., Barbieri, M., Cao, T.B.N., Varahram, N., Aljahani, A., Lidschreiber, M., Papantonis, A., Oudelaar, A.M., 2025. CTCF depletion decouples enhancer-mediated gene activation from chromatin hub formation. Nat. Struct. Mol. Biol. 10.1038/s41594-025-01555-z

48. Karr, J.P., Ferrie, J.J., Tjian, R., Darzacq, X., 2022. The transcription factor activity gradient (TAG) model: contemplating a contact-independent mechanism for enhancer–promoter communication. Genes Dev. 36, 7–16. 10.1101/gad.349160.121

49. Kawasaki, K., Fukaya, T., 2023. Functional coordination between transcription factor clustering and gene activity. Mol. Cell 83, 1605–1622.e9. 10.1016/j.molcel.2023.04.018

50. Kent, W.J., Zweig, A.S., Barber, G., Hinrichs, A.S., Karolchik, D., 2010. BigWig and BigBed: enabling browsing of large distributed datasets. Bioinformatics 26, 2204–2207. 10.1093/bioinformatics/btq351

51. Kragesteen, B.K., Spielmann, M., Paliou, C., Heinrich, V., Schöpflin, R., Esposito, A., Annunziatella, C., Bianco, S., Chiariello, A.M., Jerković, I., Harabula, I., Guckelberger, P., Pechstein, M., Wittler, L., Chan, W.-L., Franke, M., Lupiáñez, D.G., Kraft, K., Timmermann, B., Vingron, M., Visel, A., Nicodemi, M., Mundlos, S., Andrey, G., 2018. Dynamic 3D chromatin architecture contributes to enhancer specificity and limb morphogenesis. Nat. Genet. 50, 1463–1473. 10.1038/s41588-018-0221-x

52. Krebs, A.R., Imanci, D., Hoerner, L., Gaidatzis, D., Burger, L., Schübeler, D., 2017. Genome-wide Single-Molecule Footprinting Reveals High RNA Polymerase II Turnover at Paused Promoters. Mol. Cell 67, 411–422.e4. 10.1016/j.molcel.2017.06.027

53. Kubo, N., Ishii, H., Xiong, X., Bianco, S., Meitinger, F., Hu, R., Hocker, J.D., Conte, M., Gorkin, D., Yu, M., Li, B., Dixon, J.R., Hu, M., Nicodemi, M., Zhao, H., Ren, B., 2021. Promoter-proximal CTCF binding promotes distal enhancer-dependent gene activation. Nat. Struct. Mol. Biol. 28, 152–161. 10.1038/s41594-020-00539-5

54. Langmead, B., Salzberg, S.L., 2012. Fast gapped-read alignment with Bowtie 2. Nat. Methods 9, 357–359. 10.1038/nmeth.1923

55. Larke, M.S.C., Schwessinger, R., Nojima, T., Telenius, J., Beagrie, R.A., Downes, D.J., Oudelaar, A.M., Truch, J., Graham, B., Bender, M.A., Proudfoot, N.J., Higgs, D.R., Hughes, J.R., 2021. Enhancers predominantly regulate gene expression during differentiation via transcription initiation. Mol. Cell 81, 983–997.e7. 10.1016/j.molcel.2021.01.002

56. Larson, D.R., Fritzsch, C., Sun, L., Meng, X., Lawrence, D.S., Singer, R.H., 2013. Direct observation of frequency modulated transcription in single cells using light activation. eLife 2, e00750. 10.7554/eLife.00750

57. Larsson, A.J.M., Johnsson, P., Hagemann-Jensen, M., Hartmanis, L., Faridani, O.R., Reinius, B., Segerstolpe, Å., Rivera, C.M., Ren, B., Sandberg, R., 2019. Genomic encoding of transcriptional burst kinetics. Nature 565, 251–254. 10.1038/s41586-018-0836-1

58. Lee, K., Hsiung, C.C.-S., Huang, P., Raj, A., Blobel, G.A., 2015. Corrigendum: Dynamic enhancer-gene body contacts during transcription elongation. Genes Dev. 29, 2217.

59. Lettice, L.A., Horikoshi, T., Heaney, S.J.H., Van Baren, M.J., Van Der Linde, H.C., Breedveld, G.J., Joosse, M., Akarsu, N., Oostra, B.A., Endo, N., Shibata, M., Suzuki, M., Takahashi, E., Shinka, T., Nakahori, Y., Ayusawa, D., Nakabayashi, K., Scherer, S.W., Heutink, P., Hill, R.E., Noji, S., 2002. Disruption of a long-range cis-acting regulator for *Shh* causes preaxial polydactyly. Proc. Natl. Acad. Sci. 99, 7548–7553. 10.1073/pnas.112212199

60. Li, H., Handsaker, B., Wysoker, A., Fennell, T., Ruan, J., Homer, N., Marth, G., Abecasis, G., Durbin, R., 1000 Genome Project Data Processing Subgroup, 2009. The Sequence Alignment/Map format and SAMtools. Bioinformatics 25, 2078–2079. 10.1093/bioinformatics/btp352

61. Li, X., Levine, M., 2024. What are tethering elements? Curr. Opin. Genet. Dev. 84, 102151. 10.1016/j.gde.2023.102151

62. Li, X., Tang, X., Bing, X., Catalano, C., Li, T., Dolsten, G., Wu, C., Levine, M., 2023. GAGA-associated factor fosters loop formation in the Drosophila genome. Mol. Cell 83, 1519–1526.e4. 10.1016/j.molcel.2023.03.011

63. Lieberman-Aiden, E., Van Berkum, N.L., Williams, L., Imakaev, M., Ragoczy, T., Telling, A., Amit, I., Lajoie, B.R., Sabo, P.J., Dorschner, M.O., Sandstrom, R., Bernstein, B., Bender, M.A., Groudine, M., Gnirke, A., Stamatoyannopoulos, J., Mirny, L.A., Lander, E.S., Dekker, J., 2009. Comprehensive Mapping of Long-Range Interactions Reveals Folding Principles of the Human Genome. Science 326, 289–293. 10.1126/science.1181369

64. Long, H.K., Osterwalder, M., Welsh, I.C., Hansen, K., Davies, J.O.J., Liu, Y.E., Koska, M., Adams, A.T., Aho, R., Arora, N., Ikeda, K., Williams, R.M., Sauka-Spengler, T., Porteus, M.H., Mohun, T., Dickel, D.E., Swigut, T., Hughes, J.R., Higgs, D.R., Visel, A., Selleri, L., Wysocka, J., 2020. Loss of Extreme Long-Range Enhancers in Human Neural Crest Drives a Craniofacial Disorder. Cell Stem Cell 27, 765–783.e14. 10.1016/j.stem.2020.09.001

65. Long, H.K., Prescott, S.L., Wysocka, J., 2016. Ever-Changing Landscapes: Transcriptional Enhancers in Development and Evolution. Cell 167, 1170–1187. 10.1016/j.cell.2016.09.018

66. Lysakovskaia, K., Devadas, A., Schwalb, B., Lidschreiber, M., Cramer, P., 2025. Promoter-proximal RNA polymerase II termination regulates transcription during human cell type transition. Nat. Struct. Mol. Biol. 32, 995–1005. 10.1038/s41594-025-01486-9

67. Mach, P., Kos, P.I., Zhan, Y., Cramard, J., Gaudin, S., Tünnermann, J., Marchi, E., Eglinger, J., Zuin, J., Kryzhanovska, M., Smallwood, S., Gelman, L., Roth, G., Nora, E.P., Tiana, G., Giorgetti, L., 2022. Cohesin and CTCF control the dynamics of chromosome folding. Nat. Genet. 54, 1907–1918. 10.1038/s41588-022-01232-7

68. Mariner-Faulí, M., Sánchez-Gaya, V., Robert, S.M., Respuela-Alonso, P., De La Cruz-Molina, S., Lobato-Moreno, S., Trovato, M., Prummel, K.D., Noh, K.-M., Zaugg, J.B., Rada-Iglesias, Á., 2025. Dual role of ZIC2 during neural induction: from pioneer transcription factor to enhancer activator. 10.1101/2025.07.23.666365

69. Marshall, W.F., Straight, A., Marko, J.F., Swedlow, J., Dernburg, A., Belmont, A., Murray, A.W., Agard, D.A., Sedat, J.W., 1997. Interphase chromosomes undergo constrained diffusional motion in living cells. Curr. Biol. 7, 930–939. 10.1016/S0960-9822(06)00412-X

70. Mateo, L.J., Murphy, S.E., Hafner, A., Cinquini, I.S., Walker, C.A., Boettiger, A.N., 2019. Visualizing DNA folding and RNA in embryos at single-cell resolution. Nature 568, 49–54. 10.1038/s41586-019-1035-4

71. McCord, R.P., Kaplan, N., Giorgetti, L., 2020. Chromosome Conformation Capture and Beyond: Toward an Integrative View of Chromosome Structure and Function. Mol. Cell 77, 688–708. 10.1016/j.molcel.2019.12.021

72. Meeussen, J.V.W., Pomp, W., Brouwer, I., de Jonge, W.J., Patel, H.P., Lenstra, T.L., 2023. Transcription factor clusters enable target search but do not contribute to target gene activation. Nucleic Acids Res. 51, 5449–5468. 10.1093/nar/gkad227

73. Nagel, M., Taatjes, D.J., 2025. Regulation of RNA polymerase II transcription through re-initiation and bursting. Mol. Cell 85, 1907–1919. 10.1016/j.molcel.2025.04.011

74. Narita, T., Ito, S., Higashijima, Y., Chu, W.K., Neumann, K., Walter, J., Satpathy, S., Liebner, T., Hamilton, W.B., Maskey, E., Prus, G., Shibata, M., Iesmantavicius, V., Brickman, J.M., Anastassiadis, K., Koseki, H., Choudhary, C., 2021. Enhancers are activated by p300/CBP activity-dependent PIC assembly, RNAPII recruitment, and pause release. Mol. Cell 81, 2166–2182.e6. 10.1016/j.molcel.2021.03.008

75. Noonan, J.P., McCallion, A.S., 2010. Genomics of Long-Range Regulatory Elements. Annu. Rev. Genomics Hum. Genet. 11, 1–23. 10.1146/annurev-genom-082509-141651

76. Nora, E.P., Goloborodko, A., Valton, A.-L., Gibcus, J.H., Uebersohn, A., Abdennur, N., Dekker, J., Mirny, L.A., Bruneau, B.G., 2017. Targeted Degradation of CTCF Decouples Local Insulation of Chromosome Domains from Genomic Compartmentalization. Cell 169, 930–944.e22. 10.1016/j.cell.2017.05.004

77. Ong, C.-T., Corces, V.G., 2014. CTCF: an architectural protein bridging genome topology and function. Nat. Rev. Genet. 15, 234–246. 10.1038/nrg3663

78. Oudelaar, A.M., Davies, J.O.J., Hanssen, L.L.P., Telenius, J.M., Schwessinger, R., Liu, Y., Brown, J.M., Downes, D.J., Chiariello, A.M., Bianco, S., Nicodemi, M., Buckle, V.J., Dekker, J., Higgs, D.R., Hughes, J.R., 2018. Single-allele chromatin interactions identify regulatory hubs in dynamic compartmentalized domains. Nat. Genet. 50, 1744–1751. 10.1038/s41588-018-0253-2

79. Pachano, T., Sánchez-Gaya, V., Ealo, T., Mariner-Faulí, M., Bleckwehl, T., Asenjo, H.G., Respuela, P., Cruz-Molina, S., Muñoz-San Martín, M., Haro, E., Van IJcken, W.F.J., Landeira, D., Rada-Iglesias, A., 2021. Orphan CpG islands amplify poised enhancer regulatory activity and determine target gene responsiveness. Nat. Genet. 53, 1036–1049. 10.1038/s41588-021-00888-x

80. Palacio, M., Taatjes, D.J., 2022. Merging Established Mechanisms with New Insights: Condensates, Hubs, and the Regulation of RNA Polymerase II Transcription. J. Mol. Biol. 434, 167216. 10.1016/j.jmb.2021.167216

81. Perez, G., Barber, G.P., Benet-Pages, A., Casper, J., Clawson, H., Diekhans, M., Fischer, C., Gonzalez, J.N., Hinrichs, A.S., Lee, C.M., Nassar, L.R., Raney, B.J., Speir, M.L., van Baren, M.J., Vaske, C.J., Haussler, D., Kent, W.J., Haeussler, M., 2025. The UCSC Genome Browser database: 2025 update. Nucleic Acids Res. 53, D1243–D1249. 10.1093/nar/gkae974

82. Phillips-Cremins, J.E., Sauria, M.E.G., Sanyal, A., Gerasimova, T.I., Lajoie, B.R., Bell, J.S.K., Ong, C.-T., Hookway, T.A., Guo, C., Sun, Y., Bland, M.J., Wagstaff, W., Dalton, S., McDevitt, T.C., Sen, R., Dekker, J., Taylor, J., Corces, V.G., 2013. Architectural Protein Subclasses Shape 3D Organization of Genomes during Lineage Commitment. Cell 153, 1281–1295. 10.1016/j.cell.2013.04.053

83. Pollex, T., Rabinowitz, A., Gambetta, M.C., Marco-Ferreres, R., Viales, R.R., Jankowski, A., Schaub, C., Furlong, E.E.M., 2024. Enhancer–promoter interactions become more instructive in the transition from cell-fate specification to tissue differentiation. Nat. Genet. 56, 686–696. 10.1038/s41588-024-01678-x

84. Raj, A., Peskin, C.S., Tranchina, D., Vargas, D.Y., Tyagi, S., 2006. Stochastic mRNA Synthesis in Mammalian Cells. PLoS Biol. 4, e309. 10.1371/journal.pbio.0040309

85. Ramírez, F., Ryan, D.P., Grüning, B., Bhardwaj, V., Kilpert, F., Richter, A.S., Heyne, S., Dündar, F., Manke, T., 2016. deepTools2: a next generation web server for deep-sequencing data analysis. Nucleic Acids Res. 44, W160–W165. 10.1093/nar/gkw257

86. Rao, S.S.P., Huang, S.-C., Glenn St Hilaire, B., Engreitz, J.M., Perez, E.M., Kieffer-Kwon, K.- R., Sanborn, A.L., Johnstone, S.E., Bascom, G.D., Bochkov, I.D., Huang, X., Shamim, M.S., Shin, J., Turner, D., Ye, Z., Omer, A.D., Robinson, J.T., Schlick, T., Bernstein, B.E., Casellas, R., Lander, E.S., Aiden, E.L., 2017. Cohesin Loss Eliminates All Loop Domains. Cell 171, 305–320.e24. 10.1016/j.cell.2017.09.026

87. Rao, S.S.P., Huntley, M.H., Durand, N.C., Stamenova, E.K., Bochkov, I.D., Robinson, J.T., Sanborn, A.L., Machol, I., Omer, A.D., Lander, E.S., Aiden, E.L., 2014. A 3D Map of the Human Genome at Kilobase Resolution Reveals Principles of Chromatin Looping. Cell 159, 1665– 1680. 10.1016/j.cell.2014.11.021

88. Ren, G., Jin, W., Cui, K., Rodrigez, J., Hu, G., Zhang, Z., Larson, D.R., Zhao, K., 2017. CTCF-Mediated Enhancer-Promoter Interaction Is a Critical Regulator of Cell-to-Cell Variation of Gene Expression. Mol. Cell 67, 1049–1058.e6. 10.1016/j.molcel.2017.08.026

89. Rinzema, N.J., Sofiadis, K., Tjalsma, S.J.D., Verstegen, M.J.A.M., Oz, Y., Valdes-Quezada, C., Felder, A.-K., Filipovska, T., Van Der Elst, S., De Andrade Dos Ramos, Z., Han, R., Krijger, P.H.L., De Laat, W., 2022. Building regulatory landscapes reveals that an enhancer can recruit cohesin to create contact domains, engage CTCF sites and activate distant genes. Nat. Struct. Mol. Biol. 29, 563–574. 10.1038/s41594-022-00787-7

90. Sabari, B.R., Dall’Agnese, A., Boija, A., Klein, I.A., Coffey, E.L., Shrinivas, K., Abraham, B.J., Hannett, N.M., Zamudio, A.V., Manteiga, J.C., Li, C.H., Guo, Y.E., Day, D.S., Schuijers, J., Vasile, E., Malik, S., Hnisz, D., Lee, T.I., Cisse, I.I., Roeder, R.G., Sharp, P.A., Chakraborty, A.K., Young, R.A., 2018. Coactivator condensation at super-enhancers links phase separation and gene control. Science 361, eaar3958. 10.1126/science.aar3958

91. Sanborn, A.L., Rao, S.S.P., Huang, S.-C., Durand, N.C., Huntley, M.H., Jewett, A.I., Bochkov, I.D., Chinnappan, D., Cutkosky, A., Li, J., Geeting, K.P., Gnirke, A., Melnikov, A., McKenna, D., Stamenova, E.K., Lander, E.S., Aiden, E.L., 2015. Chromatin extrusion explains key features of loop and domain formation in wild-type and engineered genomes. Proc. Natl. Acad. Sci. 112. 10.1073/pnas.1518552112

92. Schwarzer, W., Abdennur, N., Goloborodko, A., Pekowska, A., Fudenberg, G., Loe-Mie, Y., Fonseca, N.A., Huber, W., Haering, C.H., Mirny, L., Spitz, F., 2017. Two independent modes of chromatin organization revealed by cohesin removal. Nature 551, 51–56. 10.1038/nature24281

93. Shimizu, R., Tanaka, M., Tsutsumi, S., Aburatani, H., Yamazaki, Y., Homme, M., Kitagawa, Y., Nakamura, T., 2018. EWS - FLI 1 regulates a transcriptional program in cooperation with Foxq1 in mouse Ewing sarcoma. Cancer Sci. 109, 2907–2918. 10.1111/cas.13710

94. Spielmann, M., Lupiáñez, D.G., Mundlos, S., 2018. Structural variation in the 3D genome. Nat. Rev. Genet. 19, 453–467. 10.1038/s41576-018-0007-0

95. Tang, Z., Luo, O.J., Li, X., Zheng, M., Zhu, J.J., Szalaj, P., Trzaskoma, P., Magalska, A., Wlodarczyk, J., Ruszczycki, B., Michalski, P., Piecuch, E., Wang, P., Wang, D., Tian, S.Z., Penrad-Mobayed, M., Sachs, L.M., Ruan, X., Wei, C.-L., Liu, E.T., Wilczynski, G.M., Plewczynski, D., Li, G., Ruan, Y., 2015. CTCF-Mediated Human 3D Genome Architecture Reveals Chromatin Topology for Transcription. Cell 163, 1611–1627. 10.1016/j.cell.2015.11.024

96. Trojanowski, J., Frank, L., Rademacher, A., Mücke, N., Grigaitis, P., Rippe, K., 2022. Transcription activation is enhanced by multivalent interactions independent of phase separation. Mol. Cell 82, 1878–1893.e10. 10.1016/j.molcel.2022.04.017

97. Tunnacliffe, E., Chubb, J.R., 2020. What Is a Transcriptional Burst? Trends Genet. 36, 288–297. 10.1016/j.tig.2020.01.003

98. Tünnermann, J., Roth, G., Cramard, J., Giorgetti, L., 2025. Enhancer control of promoter activity and variability *via* frequency modulation of clustered transcriptional bursts. 10.1101/2025.03.26.645410

99. Visel, A., Blow, M.J., Li, Z., Zhang, T., Akiyama, J.A., Holt, A., Plajzer-Frick, I., Shoukry, M., Wright, C., Chen, F., Afzal, V., Ren, B., Rubin, E.M., Pennacchio, L.A., 2009. ChIP-seq accurately predicts tissue-specific activity of enhancers. Nature 457, 854–858. 10.1038/nature07730

100. Walters, M.C., Fiering, S., Eidemiller, J., Magis, W., Groudine, M., Martin, D.I., 1995. Enhancers increase the probability but not the level of gene expression. Proc. Natl. Acad. Sci. 92, 7125–7129. 10.1073/pnas.92.15.7125

101. Wang, H., Fan, Z., Shliaha, P.V., Miele, M., Hendrickson, R.C., Jiang, X., Helin, K., 2023. H3K4me3 regulates RNA polymerase II promoter-proximal pause-release. Nature 615, 339–348. 10.1038/s41586-023-05780-8

102. Whyte, W.A., Orlando, D.A., Hnisz, D., Abraham, B.J., Lin, C.Y., Kagey, M.H., Rahl, P.B., Lee, T.I., Young, R.A., 2013. Master Transcription Factors and Mediator Establish Super-Enhancers at Key Cell Identity Genes. Cell 153, 307–319. 10.1016/j.cell.2013.03.035

103. Wissink, E.M., Vihervaara, A., Tippens, N.D., Lis, J.T., 2019. Nascent RNA analyses: tracking transcription and its regulation. Nat. Rev. Genet. 20, 705–723. 10.1038/s41576-019-0159-6

104. Wu, J., Chen, B., Liu, Y., Ma, L., Huang, W., Lin, Y., 2022. Modulating gene regulation function by chemically controlled transcription factor clustering. Nat. Commun. 13, 2663. 10.1038/s41467-022-30397-2

105. Xiao, J.Y., Hafner, A., Boettiger, A.N., 2021. How subtle changes in 3D structure can create large changes in transcription. eLife 10, e64320. 10.7554/eLife.64320

106. Yan, J., Chen, S.-A.A., Local, A., Liu, T., Qiu, Y., Dorighi, K.M., Preissl, S., Rivera, C.M., Wang, C., Ye, Z., Ge, K., Hu, M., Wysocka, J., Ren, B., 2018. Histone H3 lysine 4 monomethylation modulates long-range chromatin interactions at enhancers. Cell Res. 28, 204–220. 10.1038/cr.2018.1

107. Yanchus, C., Drucker, K.L., Kollmeyer, T.M., Tsai, R., Winick-Ng, W., Liang, M., Malik, A., Pawling, J., De Lorenzo, S.B., Ali, A., Decker, P.A., Kosel, M.L., Panda, A., Al-Zahrani, K.N., Jiang, L., Browning, J.W.L., Lowden, C., Geuenich, M., Hernandez, J.J., Gosio, J.T., Ahmed, M., Loganathan, S.K., Berman, J., Trcka, D., Michealraj, K.A., Fortin, J., Carson, B., Hollingsworth, E.W., Jacinto, S., Mazrooei, P., Zhou, L., Elia, A., Lupien, M., He, H.H., Murphy, D.J., Wang, L., Abyzov, A., Dennis, J.W., Maass, P.G., Campbell, K., Wilson, M.D., Lachance, D.H., Wrensch, M., Wiencke, J., Mak, T., Pennacchio, L.A., Dickel, D.E., Visel, A., Wrana, J., Taylor, M.D., Zadeh, G., Dirks, P., Eckel-Passow, J.E., Attisano, L., Pombo, A., Ida, C.M., Kvon, E.Z., Jenkins, R.B., Schramek, D., 2022. A noncoding single-nucleotide polymorphism at 8q24 drives *IDH1* -mutant glioma formation. Science 378, 68–78. 10.1126/science.abj2890

109. Yang, J.H., Hansen, A.S., 2024. Enhancer selectivity in space and time: from enhancer– promoter interactions to promoter activation. Nat. Rev. Mol. Cell Biol. 25, 574–591. 10.1038/s41580-024-00710-6

110. Zhang, Q., Bhattacharya, S., Andersen, M.E., 2013. Ultrasensitive response motifs: basic amplifiers in molecular signalling networks. Open Biol. 3, 130031. 10.1098/rsob.130031

111. Zug, R., 2022. Developmental disorders caused by haploinsufficiency of transcriptional regulators: a perspective based on cell fate determination. Biol. Open 11, bio058896. 10.1242/bio.058896

112. Zuin, J., Roth, G., Zhan, Y., Cramard, J., Redolfi, J., Piskadlo, E., Mach, P., Kryzhanovska, M., Tihanyi, G., Kohler, H., Eder, M., Leemans, C., Van Steensel, B., Meister, P., Smallwood, S., Giorgetti, L., 2022. Nonlinear control of transcription through enhancer–promoter interactions. Nature 604, 571–577. 10.1038/s41586-022-04570-y

